# Evidence that recurrent Group A streptococcus tonsillitis is an immunosusceptibility disease involving antibody deficiency and aberrant Tfh cells

**DOI:** 10.1101/356741

**Authors:** Jennifer M. Dan, Colin Havenar-Daughton, Kayla Kendric, Kirti Kaushik, Sandy L. Rosales, Erika Anderson, Christopher LaRock, Pandurangan Vijayanand, Gregory Seumois, David Layfield, Ramsey Cutress, Christian Ottensmier, Cecilia Lindestam Arlehamn, Alessandro Sette, Victor Nizet, Marcella Bothwell, Matthew Brigger, Shane Crotty

## Abstract

**One Sentence Summary:** Recurrent tonsillitis is a multifactorial disease associated with an aberrant tonsillar germinal center response to Group A Streptococcus.

**ABSTRACT:** Recurrent Group A Streptococcus (GAS) tonsillitis (RT) is a common indication for pediatric tonsillectomy. ‘Strep throat’ is highly prevalent among children; yet, it is unknown why some children develop RT. To gain insights into this classic childhood disease, we performed phenotypic, genotypic, and functional studies on pediatric GAS RT and non-RT tonsils. We observed significantly smaller germinal centers in GAS RT tonsils, and underrepresentation of GAS-specific germinal center follicular helper (GC Tfh) CD4^+^ T cells. RT children exhibited reduced antibody responses to GAS virulence factor SpeA. Risk and protective HLA Class II alleles for RT were identified. Finally, SpeA induced granzyme B^+^ GC Tfh cells in RT tonsils that had capacity to kill B cells. Together, these observations suggest that RT susceptibility can occur due to genetic differences that can result in aberrant GC Tfh cells and poor antibody responses to GAS SpeA.

## INTRODUCTION

‘Strep throat’ is one of the most prevalent human infections, with an estimated 600 million cases worldwide each year *(1)*. Clinical features of fever, tonsillar swelling or exudates, enlarged cervical lymph nodes, and absence of cough warrant testing for Group A Streptococcus (*S. pyogenes*, GAS) *(2, 3)*. Prompt antibiotic treatment can rapidly clear the infection *(4)*, reducing the risk of GAS-associated syndromes such as acute rheumatic fever and rheumatic heart disease *(3, 5–7)*. Some children develop recurrent tonsillitis (RT) due to GAS *(8, 9)*. Tonsillitis is a substantial healthcare burden and cause of repeated antibiotic usage. There are over 750,000 tonsillectomies performed annually in the U.S., with recurrent tonsillitis being a common indication *(2, 8, 10, 11)*. Tonsils are lymph node-like structures with open crypts designed for sampling oropharyngeal microbes. As tonsils are a nidus for GAS infection, these lymphoid tissue are anatomically poised to mount a protective immune response to the pathogen *(12, 13)*. It remains a longstanding mystery why some children get GAS RT.

To answer the question of whether some children are predisposed to RT, we established a cohort of children from the San Diego (SD) area undergoing tonsillectomies for GAS RT or for non-infectious reasons (non-RT), e.g. sleep apnea. We hypothesized that differences in the GAS-specific tonsillar immune responses may explain a predilection for some children to selectively develop GAS RT.

## RESULTS

### GC Tfh cells and GC B cells are significantly reduced in RT

RT can be a severe disease, resulting in substantial morbidity and school absences in hundreds of thousands of kids per year. By clinical history, RT children in the SD cohort had a mean of 12 tonsillitis episodes compared to 0.4 episodes among non-RT children (*P* = 0.0001, **Fig 1A**). RT and non-RT children have similar asymptomatic GAS carriage rates, ranging from 18-30% *(9, 14, 15)*, suggesting that RT is not due to differences in GAS exposure. We therefore examined possible immunological differences in children with RT. We systematically examined a range of immune cell types in tonsils from 26 RT and 39 non-RT children, ages 5-18 (**Table 1A**). Tonsils contain germinal centers, comprised of germinal center T follicular helper cells (GC Tfh), follicular dendritic cells (FDCs), and germinal center B (GC B) cells *(16)*. We observed that RT tonsils contained significantly fewer GC Tfh cells (CD4^+^ CD45RO^+^CXCR5^hi^PD-1^hi^) compared to non-RT tonsils (*P* = 0.0001, **Fig. 1B,C, fig. S1A**) with proportionately more non-Tfh (CXCR5^neg^) (*P* = 0.012, **fig. S1C**). Mantle Tfh cell (mTfh, CXCR5^int^PD-1^int^, Tfh cells outside of germinal centers) and naive CD4^+^ T cell frequencies were not significantly different (*P* = 0.076, **fig. S1B**; *P* = 0.183, **fig. S1D**). Multivariate analysis demonstrated that the GC Tfh frequencies in RT patients were highly significant with or without age as a covariate (*P* = 0.0032, **Fig. 1D**).

**Figure 1.**
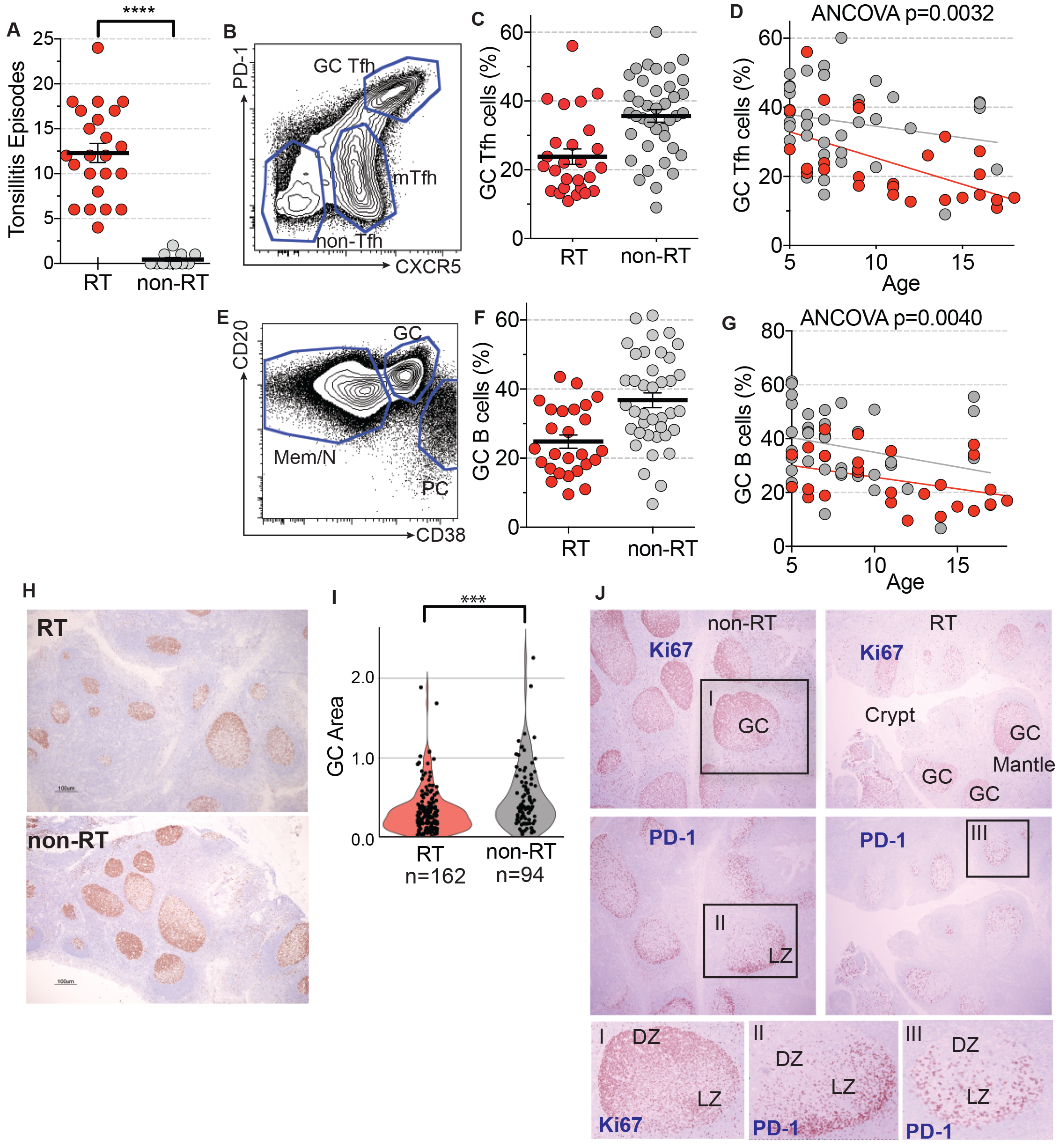
RT patients have significantly fewer GC Tfh and GC B cells. **(A)** Number of recurrent tonsillitis episodes. **(B)** Flow cytometry of GC Tfh (CXCR5^hi^PD-1^hi^CD45RO^+^CD4^+^), mTfh (CXCR5^+^PD-1^+^CD45RO^+^CD4^+^), and non-Tfh (CXCR5^−^CD45RO^+^CD4^+^) cells. **(C)** RT tonsils (n=26) have significantly fewer GC Tfh cells than non-RT tonsils (n=39). GC Tfh cells are quantified as % of total CD4^+^ T cells. **(D)** GC Tfh cells by age. **(E)** Flow cytometry of GC B cells (CD38^+^CD20^+^CD19^+^), plasma cells (CD38^hi^CD20^+^CD19^+^), and memory (CD27^hi^CD20^+^CD19^+^)/naive (CD27^−^CD20^+^CD19^+^) B cells. **(F)** GC B cells are quantified as % of total B cells. **(G)** GC B cells by age. **(H)** Representative Ki67 stained sections from RT and non-RT tonsils. **(I)** Quantitation of GC areas (μm^2^) in RT tonsils (n=21) and non-RT tonsils (n=16). Each data point represents an individual GC. **(J)** Staining of GC B cells (Ki67) and GC Tfh cells (PD-1). **** = P < 0.0001, *** = P < 0.001, ** = P < 0.01. Statistical significance determined by Mann-Whitney tests (a-c, e-f, i) and multivariate ANCOVA (d, g).

**Table 1A:**
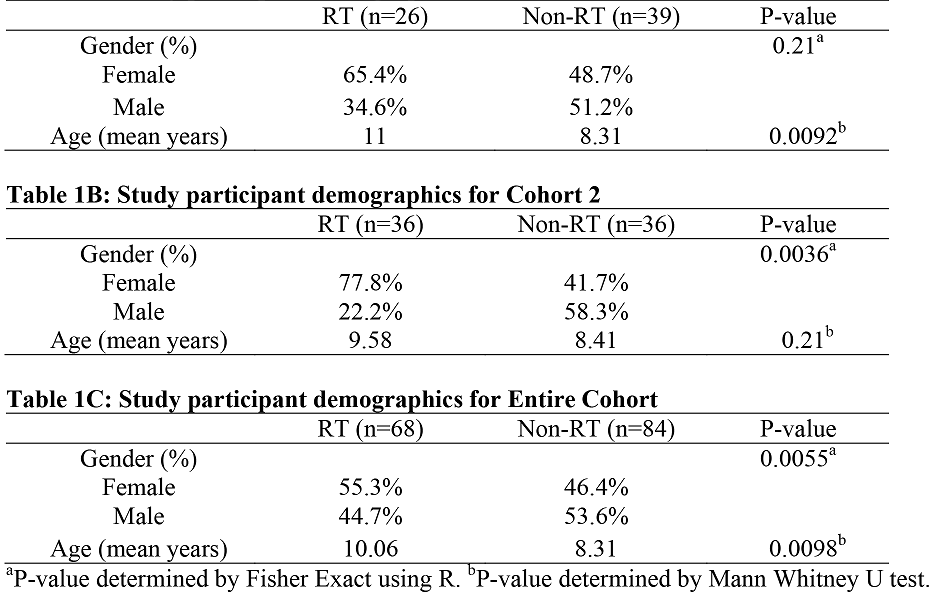
Study participant demographics for Cohort 1

Tfh cells are a distinct type of CD4^+^ T cell that provide help to B cells *(17, 18)*. Tfh cells are required for germinal centers and thus almost all affinity matured antibody responses to pathogens *(19)*. GC Tfh cells instruct the survival, proliferation, and somatic hypermutation of GC B cells. Paralleling the significant reduction in GC Tfh cells in RT patients, RT tonsils exhibited fewer GC B cells compared to non-RT tonsils (*P* = 0.0005, **Fig. 1E,F**, **fig. S1A**; *P* = 0.0040, **Fig. 1G**). GC B cells can further differentiate into plasma and memory B cells. RT tonsils had fewer plasma cells (*P* = 0.006, **fig. S1F**) but comparable memory B cell frequencies (**fig. S1A,E**). Commensurate with reduced GC B cell frequencies (*P* = 0.0002, **fig. S1A,G**), RT tonsils contained more naive B cells.

Histological examination revealed that RT tonsils had smaller germinal centers compared to non-RT tonsils (*P* < 0.002, **Fig. 1H,I**). Germinal center light and dark zones were well defined (**Fig. 1J**). Smaller germinal centers suggested a potential CD4^+^ T cell defect in RT disease, consistent with the flow cytometry data. However, differences in GC Tfh cells and germinal centers could not be directly ascribed as RT-associated or non-RT-associated without additional information (see Methods, human subjects section); thus, we explored additional parameters that establish whether the germinal center differences were associated with RT disease.

### RT disease is associated with impaired development of anti-SpeA antibodies

Diminished germinal center activity could potentially result in impaired antibody responses to GAS. Examining plasma antibodies was necessary to test this possibility; however, blood samples are not normally taken during tonsillectomies. Thus, a second cohort of patients was recruited, from whom blood samples were obtained (**Table 1B**). Antibody titers were examined against two GAS proteins: streptolysin O (SLO) — the standard GAS serodiagnostic antibody marker — and streptococcal pyrogenic exotoxin A (SpeA), a GAS virulence factor of interest. A simple expectation based on clinical history was that RT patients would have higher concentrations of GAS-specific antibodies than non-RT patients, since the former group had multiple bouts of tonsillitis (**Fig. 1A**), including experiencing a tonsillitis episode within a few months prior to surgery. However, anti-SLO IgG titers were not elevated in the RT cohort (*P* = 0.51, **Fig. 2A**). More strikingly, RT patients had significantly reduced anti-SpeA IgG titers, both when compared to non-RT patients (*P* = 0.024, **Fig. 2B**) and healthy adult volunteers (*P* = 0.0008, **Fig. 2B**). Average anti-SpeA IgG titers in RT patients were less than 10% that of healthy adult volunteers. These data indicate that impaired SpeA antibody responses are likely an attribute of RT disease, consistent with reduced germinal centers and GC Tfh deficits. SpeA antibodies have been implicated epidemiologically in immunity against severe systemic GAS infections *(20–23)*. SpeA antibodies can also be protective in a mouse GAS infection model *(24)*. Therefore, impaired production of circulating anti-SpeA IgG in RT children may be associated with RT patients’ lack of protective immunity against recurrent GAS infections.

**Figure 2.**
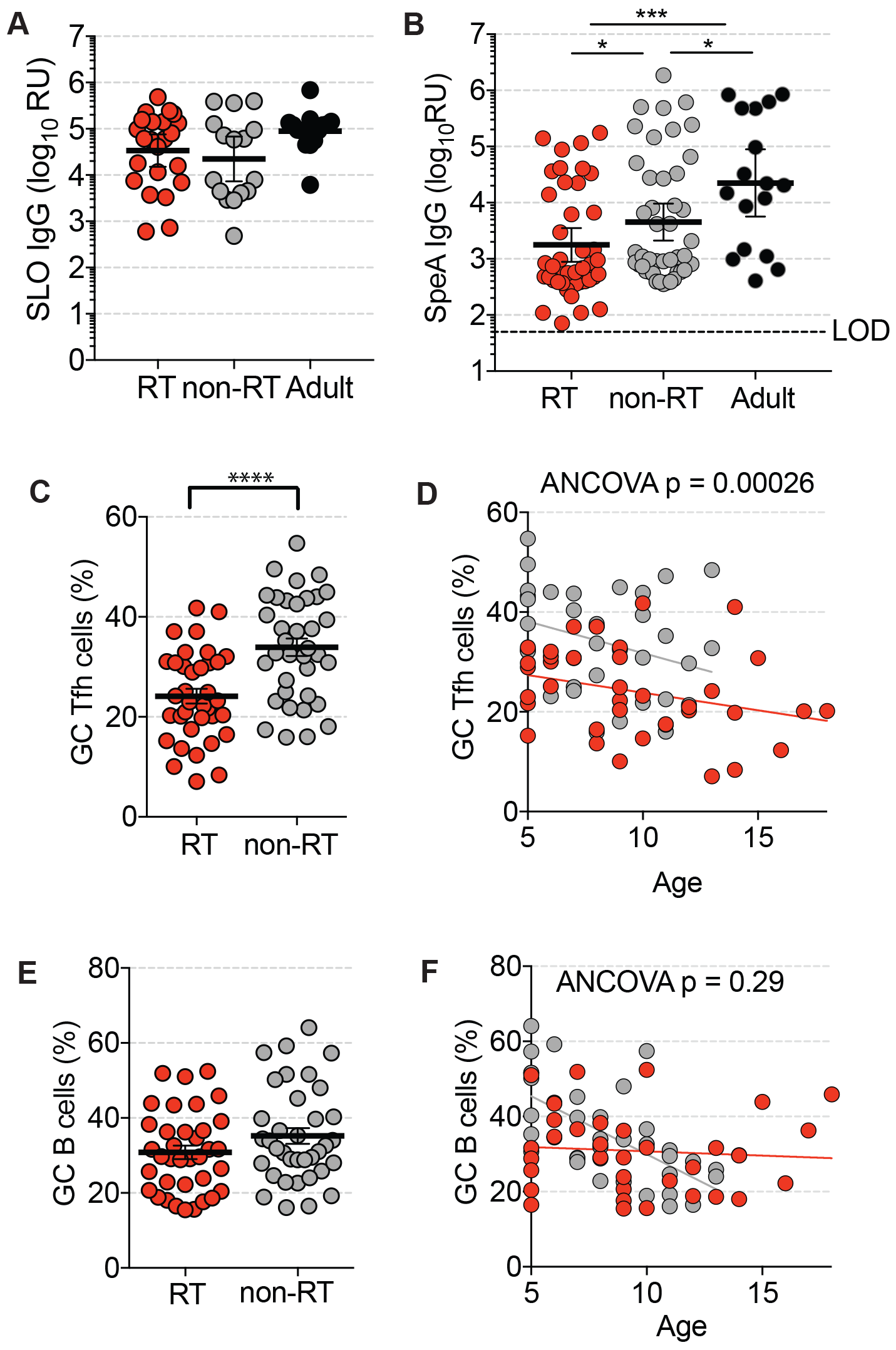
Children with RT have significantly lower titers of circulating anti-SpeA IgG. **(A)** Anti-SLO IgG titers in serum. RT children (n=23), non-RT children (n=16), and normal healthy adults (n=14). Bars indicate GMTs. **(B)** Children with RT (n=42) have significantly reduced anti-SpeA IgG compared to children with non-RT (n=45) and normal healthy adults (n=17). Bars indicate GMT, and GMTs are shown above each group. LOD = Limit of Detection **(C) 2^nd^ Cohort**: RT tonsils (n=36) have significantly fewer GC Tfh cells than non-RT tonsils (n=36). GC Tfh cells are quantified as % of total CD4^+^ T cells. **(D)** GC Tfh cells by age. **(E) 2^nd^ Cohort.** Total GC B cells are quantified as % of total B cells. P = 0.21. **(F)** GC B cells by age. * P < 0.05, *** P < 0.001. Statistical significance determined by Mann-Whitney test.

We examined tonsillar immune profiles of the second cohort. RT patients had significantly lower frequencies of GC Tfh cells than non-RT tonsils (*P* < 0.0001, **Fig. 2C**), which was independent of age (*P* = 0.00026, **Fig 2D**, **fig. S2**). Those results confirmed the observations made in the 1^st^ cohort. Significant differences in GC B cell frequencies were not observed (**Fig 2E, F**, **fig. S2**), suggesting that an RT immunological defect may be directly related to GC Tfh cells.

### GAS-specific GC Tfh cells

Phenotypic and histologic analyses of RT tonsils suggested an impairment of CD4^+^ T cell help to B cells in RT disease. We next tested for GAS-specific GC Tfh cells. Antigen-specific GC Tfh cells are difficult to identify due to their modest secretion of cytokines. We therefore developed a cytokine-independent approach to identify antigen-specific GC Tfh cells by TCR-dependent <u>A</u>ctivation <u>I</u>nduced <u>M</u>arkers (AIM) expressed upon recognition of antigen *(25, 26)*. We applied the AIM technique to quantify human GAS-specific CD4^+^ T cells (**Fig. 3A**). The non-pathogenic Gram-positive bacterium *Lactococcus lactis* was used as a negative control antigen (**Fig. 3A,B**). As children with RT experienced 12 times more tonsillitis episodes than non-RT children, a simple expectation was that RT tonsils would contain significantly more GAS-specific CD4^+^ T cells than non-RT tonsils. Instead, GAS-specific CD45RA^−^CD4^+^ T cells were not elevated in RT tonsils compared to non-RT tonsils (**Fig. 3C**). Additionally, GAS-specific CD4^+^ T cells from RT tonsils were skewed away from GC Tfh differentiation, with a lower ratio of GAS-specific GC Tfh to total GAS-specific CD4^+^ T cells (*P* = 0.025, **Fig. 3D, fig. S3B**). We observed no differences in the frequencies of GAS-specific non-Tfh cells between the groups (**fig. S3A**). Taken together, these data suggested that GAS-specific GC Tfh cell responses were disrupted in RT disease.

**Figure 3.**
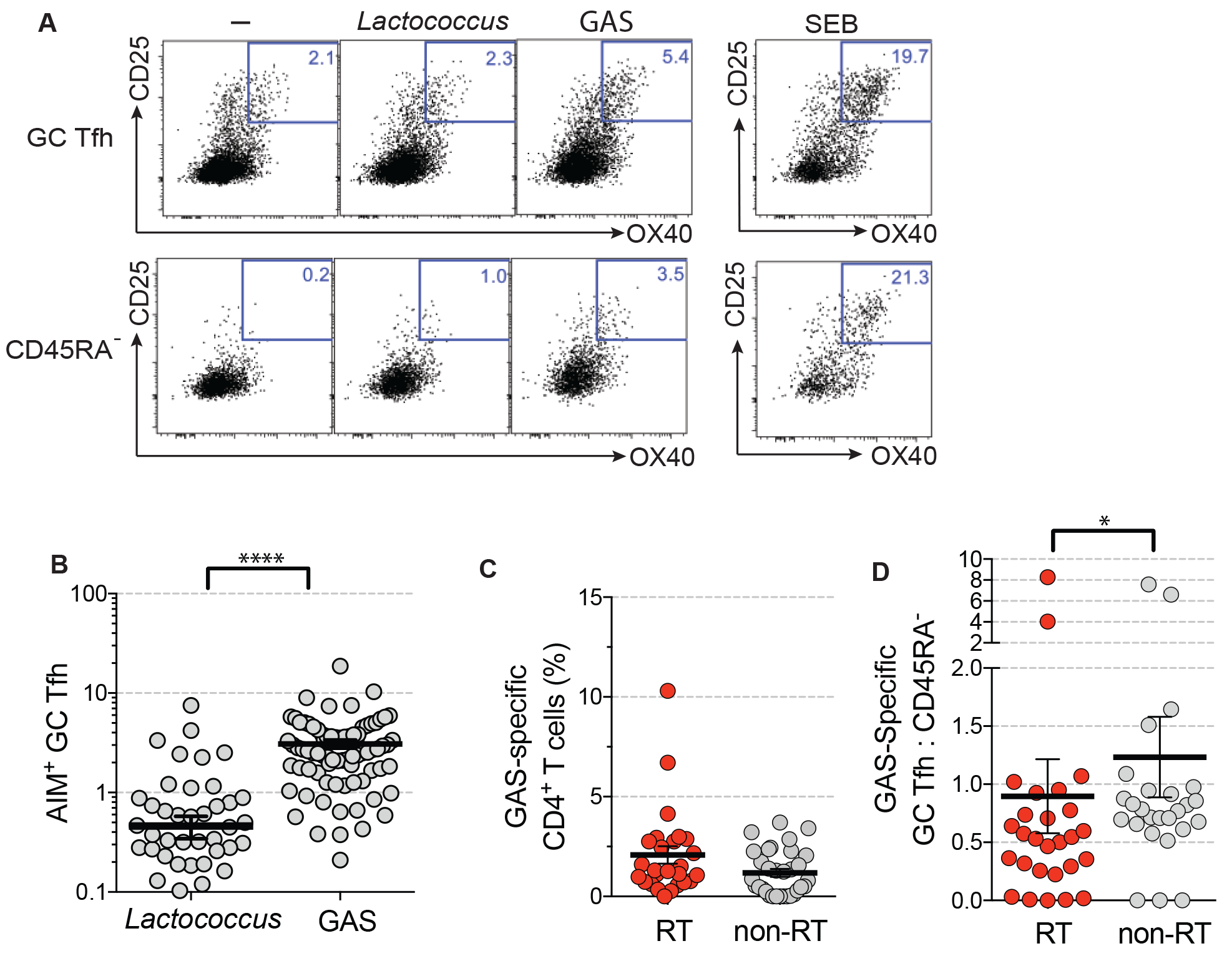
RT tonsils have reduced GAS-specific GC Tfh cells. **(A)** Flow cytometry identification of GAS-specific CD4^+^ T cells (CD45RA-) and GAS-specific GC Tfh cells (CD45RA-CXCR5^high^PD-1^high^) using an antigen-specific TCR-dependent activation induced marker (AIM) assay (OX40^+^CD25^+^, AIM25). Tonsil cells were left unstimulated or stimulated with 10μg/mL antibiotic-killed Lactococcus lactis (a non-pathogenic Gram positive bacteria which served as a negative control), 10μg/mL heat-inactivated antibiotic-killed GAS, or 1μg/mL staphylococcal enterotoxin B (SEB, positive control) for 18 hours. **(B)** Significantly more GAS-specific GC Tfh cells are detected by AIM_25_ (CD25^+^OX40^+^) compared to a negative control antigen. Cells were stimulated with heat-inactivated, antibiotic-killed GAS (‘GAS’) or antibiotic-killed *L. lactis* (‘*Lactococcus*’). **(C)** GAS-specific CD4^+^ T cell frequencies in RT and non-RT tonsils. Total antigen-experienced cells were gated (CD45RA^−^). **(D)** RT patients exhibit a bias against GAS-specific GC Tfh differentiation. Among total GAS-specific CD4^+^ T cells (AIM_25_^+^ CD45RA^−^), a smaller fraction of cells are GC Tfh cells (CXCR5^high^PD-1^high^) in RT donors compared to non-RT donors.

### RT disease is associated with HLA Class II alleles

Essentially all children are exposed to GAS during childhood *(27)*. Among patients enrolled in this study, children with RT were likely to have a significant family history of tonsillectomy (*P* = 0.0004, **Fig. 4A**). This suggested a potential genetic predisposition. Germinal center responses depend on HLA Class II antigen presentation by B cells to GC Tfh cells. Susceptibility to toxic shock syndrome and invasive forms of GAS infection have been inversely associated with HLA DQB1*06:02 *(28)*. DQB1*06:02 has also been associated with protection from the development of rheumatic heart disease *(29, 30)*, the most severe sequela of long-term untreated GAS RT and the leading cause of heart failure in children worldwide *(1, 31)*. We performed HLA typing of the entire cohort and compared the HLA allelic frequencies to ethnically matched healthy adults from the local general population (**fig. S4A**). HLA DQB1*06:02 was significantly less frequent in children with RT than in the general population (*P* = 0.042) and the combined groups of non-RT children + general population (*P* = 0.048, **Fig. 4B, fig. S4B**). No HLA DQB1*06:02 allelic frequency difference was present between non-RT patients and general population controls (*P* = 0.89) (**Fig. 4B**, fig. S4B). Overall, these data suggest that HLA allele DQB1*0602 is a protective allele from RT disease.

**Figure 4.**
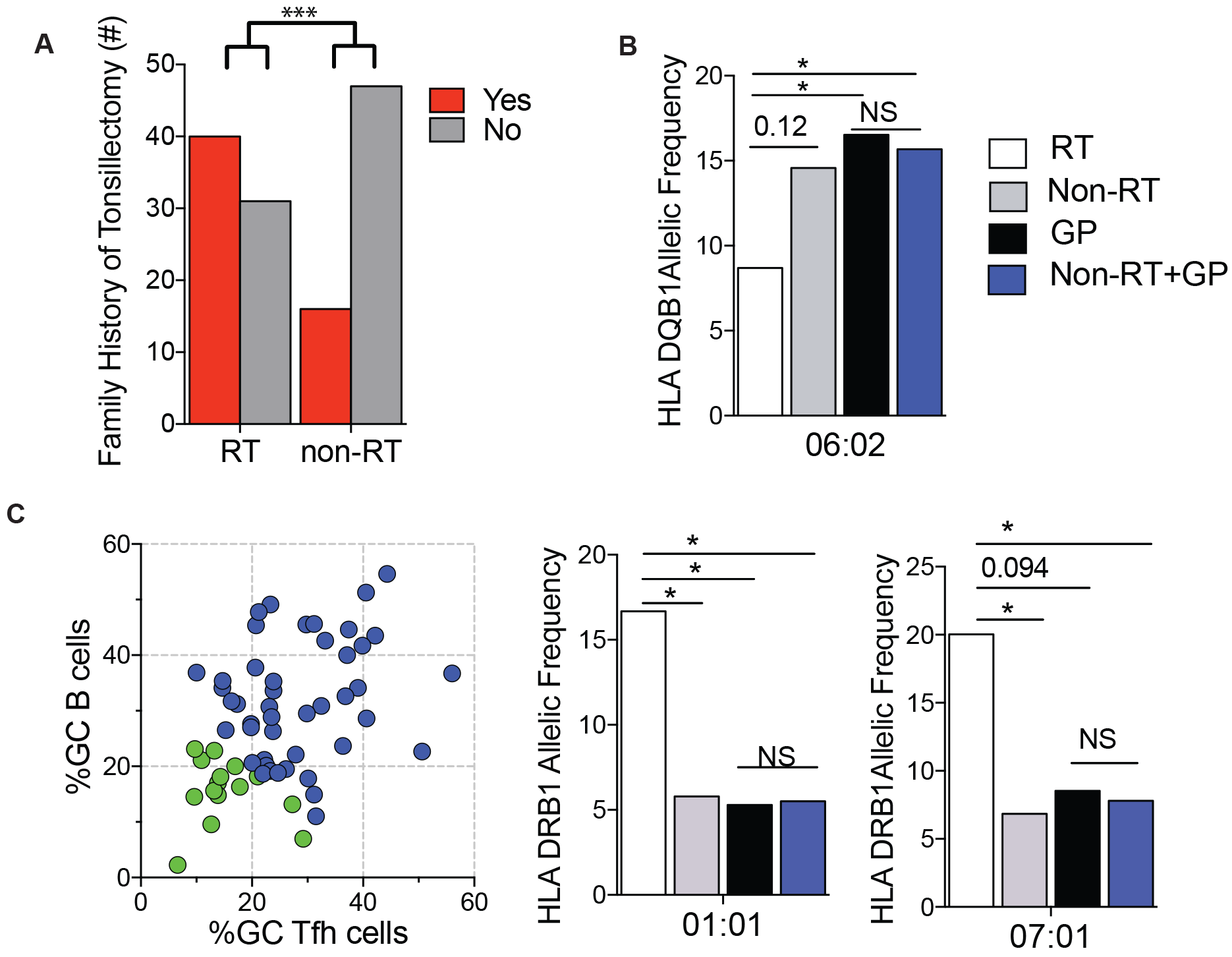
HLA associations identified in RT patients. **(A)** Family history of tonsillectomy. A significantly greater proportion of RT children have a family history of tonsillectomy than non-RT children. (RT = 71, non-RT = 63). **(B)** HLA DQB1*06:02 alleles were present at a higher frequency in non-RT patients (grey bar, n=192) compared to RT patients (white bar, n=138). HLA DQB1*06:02 alleles were present at a significantly higher frequency in the general population (black bar, n=242) and non-RT patients + general population (blue bar, n=434) compared to RT patients. **(C)** RT patients with the lowest quartile of germinal center activity, defined as lowest combined frequencies of GC Tfh and GC B cells (left panel. Green dots, n=15. GC^lo^), have a significantly higher frequency of HLA DRB1*01:01 and HLA DRB1*07:01 alleles compared to non-RT tonsils (grey bar, n=190), general population (black bar, n=246), and general population + non-RT tonsils (blue bar, n=436). RT patients HLA allele counts (white bar, n=30).
*** P < 0.001, * P < 0.05. Statistical significance determined by Fisher Exact test (a-c).

HLA alleles DRB1*01:01 *(30, 32)* and DRB1*07:01 have been linked to increased risk for rheumatic heart disease. No significant DRB1*01:01 and DRB1*07:01 allelic associations were observed among all patients enrolled in this study (**fig. S4B**). However, given that RT is a multifactorial disease, we considered that a genetic association with disease susceptibility may be more evident in RT patients exhibiting the largest germinal center deficits. HLA allelic frequencies were thus examined among children with RT with the lowest quartile of GC Tfh and GC B cells (**Fig. 4C**, **fig. S4B**, GC^lo^). These children had significantly higher frequencies of HLA DRB1*01:01 compared to the general population (*P* = 0.03), non-RT children (*P* = 0.049), and the combined control groups (*P* = 0.03, **Fig. 4C**, **fig. S4B**). Frequencies of HLA DRB1*07:01 were also elevated compare to non-RT patients and the combined control groups (*P* = 0.03, *P* = 0.03. **Fig. 4C, fig. S4B**). In contrast, no differences were identified between the non-RT and general population cohort for HLA DRB1*01:01 (*P* = 0.85) or HLA DRB1*07:01 (*P* = 0.74, **Fig. 4C**, **fig. S4B**). These data indicate that HLA DRB1*01:01 and DRB1*07:01 are risk alleles for RT. Overall, integration of HLA typing and immunophenotyping revealed relationships between RT disease, GAS, and germinal center responses.

### RT-associated HLA alleles differentially impact CD4^+^ T cell responses to GAS and the GAS superantigen SpeA

SpeA superantigen is an important GAS virulence factor. Comparison of CD4^+^ T cell reactivity using an antibiotic-killed wild-type (WT) GAS strain with or without heat inactivation, or an antibiotic-killed isogenic SpeA-deficient mutant GAS strain (*ΔspeA*), demonstrated that SpeA superantigen-mediated stimulation of CD4^+^ T cells constituted a major fraction of CD4^+^ T cell reactivity to GAS (*P* = 0.002, **Fig. 5A**, **fig. S5A**). SpeA has provided GAS with an evolutionary advantage *(24, 33, 34)*, associated with the global persistence and dominance of the M1 serotype among throat cultures. GC Tfh cells from ‘at risk’ RT tonsils were less responsive to SpeA stimulation than non-RT tonsils bearing the ‘protective’ HLA allele (*P* = 0.052, **Fig. 5B**, **fig. S5B**). While not reaching statistically significance, in light of the small N value we found the results intriguing enough to examine SpeA interactions with human CD4^+^ T cells in greater detail.

**Figure 5.**
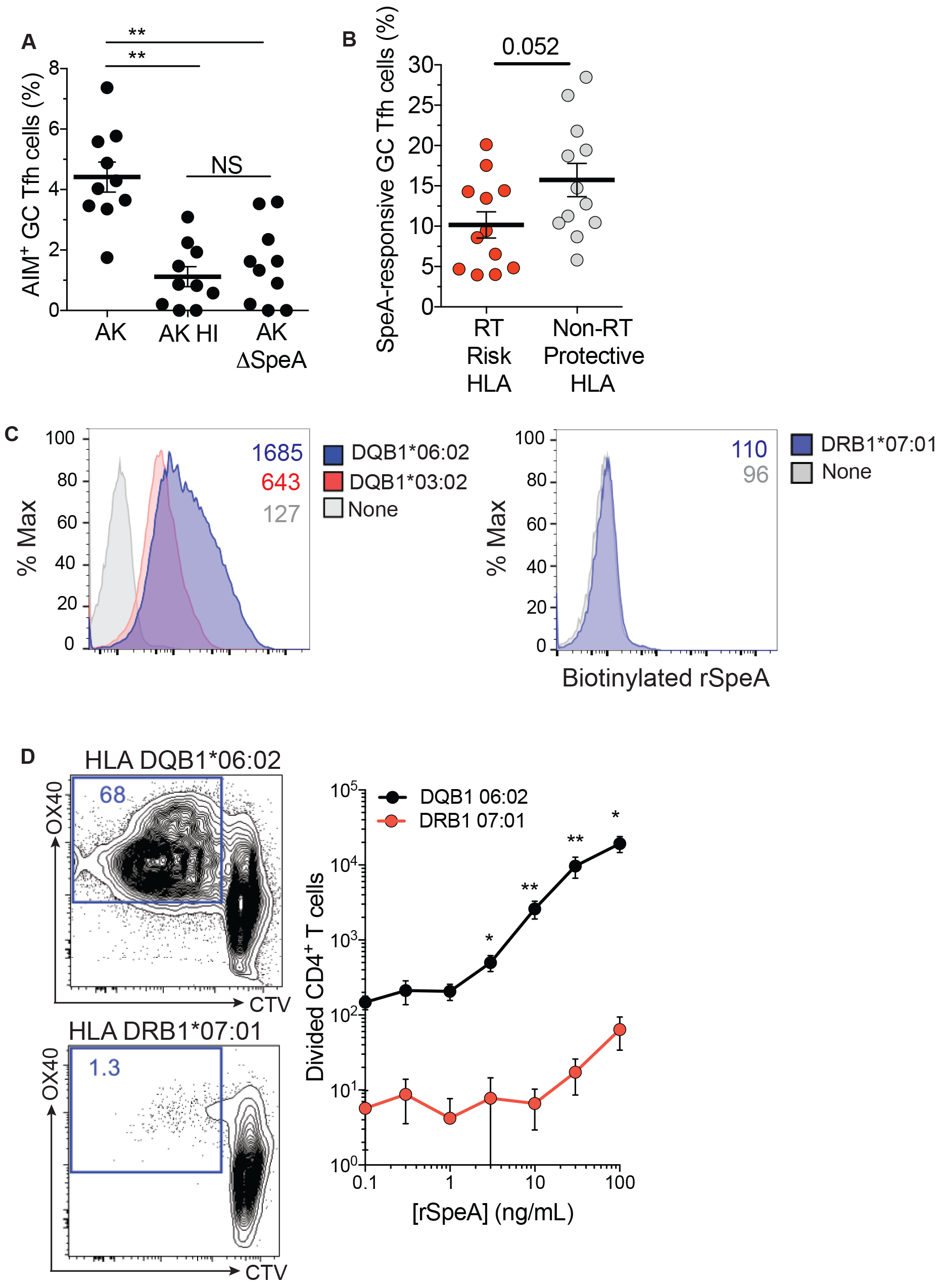
HLA associations identified in RT and non-RT patients segregate based on preferential GAS superantigen SpeA binding. **(A)** Comparison of activated (AIM^+^) GC Tfh cells following stimulation with either 10 μg/mL antibiotic-killed GAS (AK), 10 μg/mL antibiotic-killed, heat-inactivated GAS (AK HI), or 10 μg/mL antibiotic-killed SpeA deficient GAS (AK ±SpeA), n=10. **(B)** RT patients with the ‘At Risk’ HLA (n=12) have fewer SpeA-responsive GC Tfh cells compared to non-RT patients with the ‘Protective’ HLA (n=12), P=0.052. Tonsils cells were stimulated with 1μg/mL SpeA for 18 hours and background subtracted from unstimulated cells. **(C)** Histogram flow cytometric quantitation of SpeA binding. Biotinylated SpeA binds preferentially to HLA DQB1*06:02 > DQB1*03:02 > DRB1*07:01, N=3 experiments. **(D)** Magnetically sorted total CD4^+^ T cells from PBMCs of HLA DQB1*06:02^+^ donors, cocultured with SpeA and a cell line expressing HLA DQB1*06:02 proliferated significantly more compared to CD4^+^ T cells from PBMCs of HLA DRB1*07:01^+^ donors, co-cultured with SpeA (recombinant SpeA, rSpeA) and a cell line expressing HLA DRB1*07:01. N=4 experiments. ** P < 0.01, * P < 0.05 (e). Statistical significance determined by Mann Whitney test.

Mechanistic relationships between HLA class II alleles and GAS disease manifestations are unclear *(31)*, but a potential role has been suggested for SpeA *(28, 35, 36)*. We tested binding of SpeA to 19 well-defined single-allele HLA class II expressing cell lines. The highest affinity binding interaction was between SpeA and HLA DQB1*06:02 (**Fig. 5C, fig. S5C**), while moderate binding was observed to cells expressing another DQ allele, DQB1*03:01. Rapid and robust proliferation of HLA DQB1*06:02^+^ CD4^+^ T cells was observed in the presence of the superantigen (*P* = 0.0079, **Fig. 5D**, **fig. S5D**). In contrast, minimal proliferation was observed for HLA DQB1*06:02^−^ CD4^+^ T cells, including HLA DRB1*01:01^+^ or DRB1*07:01^+^ cells (**Fig. 5D** and data not shown) with minimal cell death (**fig. S5E**). In summary, highest affinity interaction of SpeA with HLA DQB1*06:02 (**Fig. 5C,D**) resulted in the most sensitive CD4^+^ T cell proliferation and was associated with reduced risk for RT disease (**Fig. 4B**).

### Granzyme B^+^ perforin^+^ GC Tfh cells are found in RT disease

While frequencies of GC Tfh cells were different between RT and non-RT patients (**Fig 1C,D**. **Fig 2C,D**), BCL6 expression by GC Tfh cells was equivalent on a per cell basis (**fig. S1H**). To identify CD4^+^ T cell factors potentially involved in SpeA superantigen-associated germinal center abnormalities in RT disease, we performed RNA sequencing (RNA-seq) on GC Tfh cells from RT and non-RT tonsils in the presence of SpeA. Strikingly, *GZMB* mRNA, encoding the cytotoxic effector protein granzyme B (GzmB), was upregulated in RT GC Tfh cells (*P* < 0.0079, **Fig. 6A**, fig. S5A, table S1, fig. S6A). GzmB is secreted by cytotoxic CD8^+^ T cells and NK cells for killing of target cells. Expression of GzmB by GC Tfh cells is contrary to the B cell help function of GC Tfh cells. Aberrant GzmB expression by GC Tfh cells could result in conversion of a GC Tfh cell from one that helps GC B cells to one that kills GC B cells, which would be a potential mechanism by which GAS could disrupt antibody responses.

**Figure 6.**
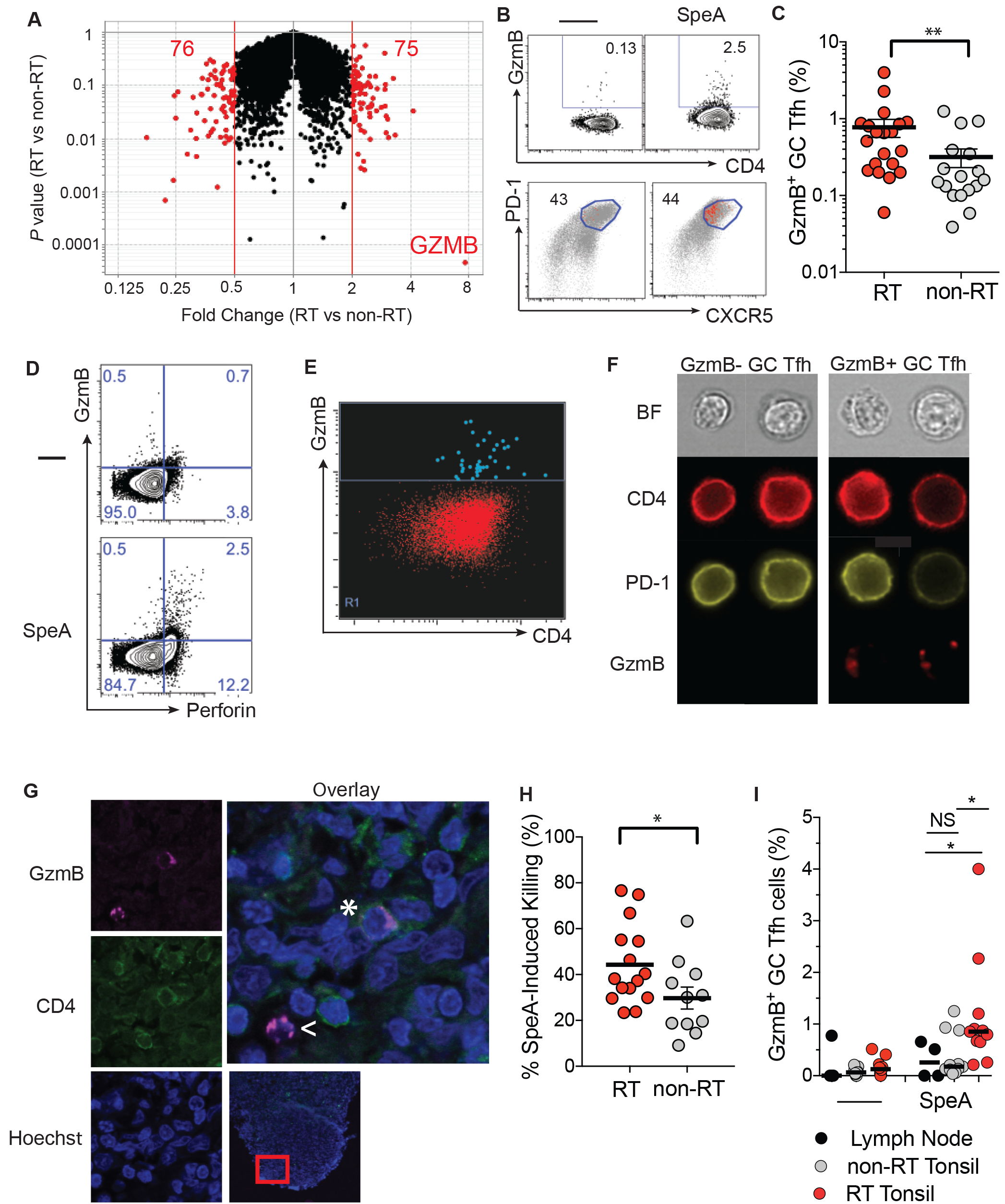
SpeA stimulation of GC Tfh cells from RT tonsils induces granzyme B and perforin. **(A)** Volcano plot showing fold change of genes in SpeA-stimulated GC Tfh cells from RT tonsils (n=5) compared to SpeA-stimulated GC Tfh cells from non-RT tonsils (n=5). Red dots denote genes with a < 2 or 2 > fold change. 76 genes exhibited a < 2-fold change and *P* < 0.1, and 75 genes with > 2-fold change and *P* < 0.1. **(B)** (top) Intracellular granzyme B expression (%) by GC Tfh cells by flow cytometry. Tonsil cells were stimulated with 1μg/mL SpeA for 24 hours. (bottom) Backgating of the granzyme B^+^ GC Tfh cells among total CD45RA^−^ CD4^+^ T cells. **(C)** SpeA stimulation revealed significantly more granzyme B^+^ GC Tfh cells in RT tonsils (n=20) compared to non-RT tonsils (n=17). **(D)** SpeA stimulated GC Tfh cells co-expressed granzyme B and perforin. FACS sorted GC Tfh cells and autologous B cells were cultured +/− SpeA for 5 days and stained for granzyme B and perforin expression. N=3 donors. **(E)** ImageStream cytometry plot of granzyme B^+^ GC Tfh cells following SpeA stimulation. GC Tfh cells were gated as CXCR5^hi^PD-1^hi^ of live CD45RA^−^CD4^+^ T cells. **(F)** ImageStream imaging of GC Tfh cells following SpeA stimulation, showing representative granzyme B^−^ and granzyme B^+^ cells. **(G)** Confocal microscopy of a granzyme B^+^ CD4^+^ T cell in a germinal center in an RT tonsil (*). A granzyme B^+^ CD8^+^ T cell is also shown for reference (<). **(H)** SpeA-stimulated GC Tfh cells are able to kill B cells. GC Tfh cells (CXCR5^hi^PD-1^hi^CD45RA^−^ CD4^+^) were co-cultured with autologous CTV-labeled B cells (CD19^+^CD38^−^). Killing was quantified as outlined in the Methods, with controls shown in **Supplementary Figure 5g-k**. N=15 RT and 11 non-RT donors. **(I)** Granzyme B expression (%) by GC Tfh cells from healthy lymph nodes and RT and non-RT tonsils. SpeA-stimulated GC Tfh cells from RT tonsils (n=11) expressed more granzyme B compared to SpeA-stimulated GC Tfh cells from non-RT tonsils (n=11) or healthy lymph nodes (n=4).
** P < 0.01, * P < 0.05. Statistical significance determined by Mann-Whitney test (c, h, i).

To determine if RT GC Tfh cells were capable of GzmB protein expression, four independent approaches were used: (1) flow cytometry of intracellular stained SpeA-stimulated GC Tfh cells, (2) ImageStream imaging cytometry of SpeA-stimulated GC Tfh cells, (3) immunofluorescence microscopy of human tonsillar tissue, and (4) killing of target cells. GC Tfh cell intracellular protein staining confirmed GzmB expression specifically induced by GAS SpeA stimulation (*P* = 0.009, **Fig. 6B-D**). Perforin expression was also induced by SpeA stimulation (**Fig. 6D**). Consistent with these findings, punctate cytoplasmic GzmB was observed in SpeA stimulated GC Tfh cells from an RT patient by ImageStream (**Fig. 6E,F**). These changes were specific to GC Tfh cells, as there were no differences in the frequencies of GzmB^+^ mTfh (**fig. S7A,D**), non-Tfh (**fig. S7B,D**) or CD8^+^ T cells (**fig. S7C,E**) between RT and non-RT donors. These GzmB^+^ GC Tfh cells were not Tregs (**fig. S7F**). GzmB was also observed histologically in some GC Tfh cells (**Fig. 6G**).

We assessed whether SpeA-stimulated GC Tfh cells were capable of killing B cells. Killing by cytotoxic CD4^+^ T cells is difficult to demonstrate *in vitro*; nevertheless killing of B cells by GC Tfh cells was observed in the presence of SpeA (**Fig. 6H**, **fig. S7G,H**). This killing was more profound with cells from RT tonsils compared to non-RT tonsils. Bystander cell death was not observed (**fig. S7I**). PHA did not stimulate GzmB expression (**fig. S7L**). B cell killing by RT GC Tfh cells in the presence of GAS SpeA was independent of Fas and FasL (**fig. S7J,K**), and was associated with perforin expression by the GzmB^+^ GC Tfh cells (**Fig. 6D**).

Lastly, we assessed whether GzmB^+^ GC Tfh cells were unique to RT. GC Tfh cells from healthy lymph nodes (LN) from patients undergoing a staging LN biopsy were compared to RT and non-RT tonsils. GzmB^+^ GC Tfh cells were sporadically detected in healthy LNs. Significantly more GzmB^+^ GC Tfh cells were observed in RT tonsils than healthy LNs, and GzmB expression was specifically induced upon SpeA stimulation (*P* = 0.025, **Fig. 6I**). GC Tfh cells from non-RT tonsils and healthy LNs were indistinguishable (**Fig. 6I**). Collectively these data suggest that SpeA is capable of deviating GC Tfh cells into GzmB^+^ killer Tfh cells and these killer Tfh cells are a pathological feature of RT disease.

## DISCUSSION

By integrating immune profiling and clinical data with transcriptomic and cell function analyses, we revealed immunologic features of GAS recurrent tonsillitis that provide evidence that RT is an immunosusceptibility disease. We observed that 1) RT patients exhibit significant reductions in GC Tfh cell frequencies; 2) RT children have impaired anti-SpeA antibody responses, which have been associated with protective immunity to GAS; 3) there are HLA Class II allele associations with RT; and 4) SpeA can induce GzmB expression in GC Tfh cells. While RT is surely a multifactorial disease, these findings indicate that the sporadic nature of recurrent tonsillitis is linked to HLA-associated genetic susceptibility differences that are mechanistically associated with differential binding of the GAS superantigen SpeA to allelic variants of human MHC II molecules. SpeA perturbation of GC Tfh cells and killing of GC B cells is a parsimonious model to explain key immunological and pathological aspects of RT.

It has been a long-standing mystery why some children get recurrent strep throat. Specific strains of GAS have been proposed as a cause of RT *(37–39)*. However, RT children and non-RT children have similar asymptomatic GAS carriage rates *(9, 14, 15)*, indicating that RT is not due to differences in GAS exposure. Our cohort consisted of children enrolled from the same geographic area to help control for differences in GAS serotypes. Globally, GAS disease burden is high and, in recent decades, the M1 serotype has remained one of the dominant strains *(40, 41)*. In this regard, it is notable that the M1 serotype possesses a bacteriophage encoding SpeA, and the acquisition of SpeA has been implicated in the dominance of the M1 pandemic strain in the United States *(42, 43)*. Here, we observed that SpeA was the dominant GAS superantigen activity on GC Tfh cells. GAS appears to take advantage of HLA class II differences via the virulence factor SpeA, rendering individuals more susceptible to pharyngitis and less likely to develop protective anti-SpeA antibody immunity from reinfection due to disruption of GC Tfh cells. SpeA skews GC Tfh cell function, resulting in cytolytic GC Tfh cells. This represents a novel immune evasion mechanism of a pathogen. RT tonsils had a higher frequency of GzmB^+^ GC Tfh cells and reduced anti-SpeA antibody responses. It is reasonable to propose that a small amount of GzmB expression may have devastating effects within the confines of a germinal center. We contend that the cytotoxicity scenario is fundamentally different for GzmB^+^ Tfh disruption of GCs than it is for CTL control of viral infection. GC B cells are probably the most pro-apoptotic cells in the body. Each GC B cell requires stimulation by a Tfh cell every few hours or it will die. Additionally, unlike most cell types, the GC B cells are all confined to a densely packed space, the germinal center. Equally importantly, by intravital microscopy, Tfh cells are constantly making short (~5 min) cognate interactions with GC B cells. Thus, in a 24 hour period, ten GC Tfh cells can make cognate interactions with 2,880 GC B cells, and an average germinal center contains only ~1,000 total GC B cells. In contrast, CTL killing of virally-infected cells takes much longer cognate interactions, with more resistant cells, over much greater 3D space. Hence, we consider it a reasonable model that it may take very little GzmB to kill a GC B cell and that GzmB^+^ Tfh could serially poison many GC B cells.

Our finding of SpeA-induced GzmB^+^perforin^+^ GC Tfh cells within tonsils also highlights the plasticity of Tfh cells. Granzyme A expressing GC Tfh cells have been described recently in human lymph nodes and tonsils *(44, 45)*. In this study, we observed no difference in the RNA expression of STAT3, granzyme A, CD57, and CRTAM between RT and non-RT tonsils (**fig. S6B-E**). However, we did observe similarities between granzyme B^+^ GC Tfh and recent reports of CD8^+^ T cells acquiring Tfh phenotypic features *(46)*. CXCR5^+^ CD8 T cells have been identified in the context of HIV, SIV, and LCMV chronic infections and have the capacity to migrate into B cell follicles and exhibit cytotoxicity *(47–50)*. Intriguingly, anti-PD1 immunotherapy predominantly rescues exhausted CD8^+^ T cells via outgrowth of CXCR5^+^ CD8^+^ T cells *(50)*. Development of CXCR5^+^ CD8^+^ T cells is associated with upregulation of known key regulators of Tfh differentiation TCF1 and BCL6 *(51)*, and a substantial reduction in the expression of GzmB by the CXCR5^+^ CD8^+^ T cells *(50, 52)*. In this study of RT GC Tfh cells, the opposite was observed; downregulation of *TCF1* and its homolog *LEF1* occurred in SpeA stimulated GC Tfh cells commensurate with GzmB upregulation, suggesting that the TCF1/LEF1 axis may be required for separation of Tfh and CTL transcriptional programs in both CD4^+^ and CD8^+^ T cells. Altogether, the data from this study suggest that conversion of GC Tfh to GzmB^+^perforin^+^ Tfh cells represents a reciprocal process to the recently described conversion of CXCR5^−^ GzmB^hi^ CD8^+^ T cells to CXCR5^+^ GzmB^lo^ CD8^+^ T cells.

This study also identified risk and protective alleles for GAS recurrent tonsillitis, and these alleles have previously been implicated in other clinical presentations of GAS infection. RT disease is associated with a lower frequency of HLA alleles observed to be protective against GAS invasive infection and toxic shock syndrome, and a higher frequency of HLA risk alleles shared with severe autoimmune rheumatic heart disease. Screening for these HLA alleles in pediatric patients with strep throat may provide a valuable prognostic indicator of the likelihood of future GAS reinfections and the utility of tonsillectomy.

In a murine HLA class II model of GAS infection, establishment of GAS infection was dependent on SpeA, and immunization with an SpeA toxoid elicited anti-SpeA IgG that was protective against GAS infection *(24, 33)*. Our data indicate that differential binding of SpeA to HLA class II alleles may predict susceptibility of individuals to GAS infection, as children with the DQB1*06:02 allele are more resistant to RT. More broadly, these data support central roles for SpeA and anti-SpeA IgG in tonsillitis pathogenesis and GAS protective immunity, respectively. An understanding of this immune evasion strategy may now allow for rational design of countermeasures. These findings indicate that an inactivated SpeA toxoid vaccine may be a simple and reasonable candidate for consideration as a strep throat and RT vaccine, as a means to reduce costly RT antibiotics treatments and surgeries per year and reduce childhood strep throat disease burden generally.

In conclusion, we provide evidence that recurrent tonsillitis is a genetic immunosusceptibility disease with a role for SpeA and GC Tfh cells. We have identified correlates of disease both on the side of the pathogen and on the side of the immune system. These findings have several implications, including the plausibility of SpeA as a potential vaccine target for RT and strep throat generally. Finally, the finding of GzmB^+^perforin^+^ GC Tfh cells points to a pathological mechanism of germinal center control.

## METHODS

### Human subjects research

Fresh tonsils were obtained from pediatric donors undergoing tonsillectomy at Rady Children’s Hospital or the Naval Medical Center. Specimens were collected at the time of surgery, at least 6 weeks after the last episode of tonsillitis, with most cases substantially further from the last episode of tonsillitis and antibiotic treatment. Beginning with later donors enrolled, at the time of tonsillectomy a blood specimen was also acquired. Informed consent was obtained from all donors under protocols approved by the institutional review boards (IRBs) of the University of California, San Diego, the La Jolla Institute for Allergy and Immunology (LJI), and the Naval Medical Center. In this study, we recruited children from the same geographic area to control for circulating strains within the community. Cohort 1was enrolled at the Naval Medical Center and Rady Children’s Hospital. Cohort 2 was enrolled only at Rady Children’s Hospital. Cohort 2 included a blood specimen on which serologies were run.

### A note on tissue sample acquisition

Tonsils are never removed from healthy children. Partial tonsil biopsies are not possible because of the small risk of life-threatening oropharnygeal hemorrhage. Cadaveric tonsils are not acceptable for research purposes, due to the highly apoptotic nature of GC B cells. Pediatric whole body organ donors are extremely rare, and those with tonsils harvested are even rarer; and those donors are regularly treated with high dose steroids and intravenous antibiotics continuously up to the moment of organ harvest, which are expected to substantially modify tonsillar biology and immune cells and thus are unaccepted for immunological comparisons.

Fresh lymph nodes were acquired from patients undergoing staging sentinel lymph node biopsy for early-stage breast cancer at University Hospital Southampton, UK, in whom said staging demonstrated the absence of lymphatic metastasis. All patients had provided informed consent for tissue donation for the purpose of clinical research study (UKCRN ID: 11947) according to protocols approved by the National Research Ethics Service following regional ethics committee review (South Central England).

### Cell processing

Tonsillar mononuclear cells were obtained by homogenizing the tissue using a wire mesh, passage through a cell strainer, and isolation via Ficoll density gradient using Histopaque 1077. Peripheral blood mononuclear cells (PBMCs) were isolated by density gradient centrifugation using Histopaque 1077 (Sigma). For PBMCs, plasma was saved after density gradient centrifugation. Cells were then washed and suspended in fetal bovine serum (FBS) containing 10% dimethyl sulfoxide, and cryopreserved in liquid nitrogen.

Single cell suspensions of lymph node-derived cells were obtained from freshly excised axillary nodes following enzymatic digest (0.15 Wünsch units/ml Liberase DL (Roche), 800 Kunitz units/ml DNAse 1 (Sigma)) over 1 hour at 37°C followed by passage through a wire mesh and 70μm cell strainer (BD Falcon). Cells were suspended in complete RPMI 1640 (Gibco + 25 mM HEPES (Sigma), Penicillin/Streptomycin (Sigma), L-Glutamine (Sigma), sodium pyruvate (Gibco) – “cRPMI”) and cryopreserved (50% decomplemented human Ab serum (Sigma), 10% Dimethyl Sulfoxide (Sigma)) in liquid nitrogen until use.

### Antibodies and Flow cytometry

Cells were labeled with fixable viability dye eFluor 780 (Thermo Fisher Scientific). FACS staining buffer consisted of 0.5% Bovine serum albumin (BSA) in phosphate buffered saline (PBS). Primary stains for leukocyte phenotyping (**Fig. 1A**) was done using fresh cells. Anti-human antibodies for surface staining of fresh tonsils are listed here, by company, Thermo Fisher Scientific: CD19 e780 (clone HIB19), CD14 e780 (clone 61D3), CD16 e780 (clone eBioCB16), CD3 e780 (clone UCHT1), CD25 PE-Cyanine 7 (clone BC96), PD-1 PE (clone eBioJ105), CD38 PE-cyanine 7 (clone HIT2), ICOS PerCP-eFluor710 (clone ISA-3), CD27 PerCP-eFluor710 (clone O323), CD45RO FITC (clone UCHL1); Biolegend: CD20 BV570 (clone 2H7), CD19 AF700 (clone HIB19), CXCR5 BV421 (clone J252D4); BD Biosciences CD3 AF700 (clone UCHT1) and CD4 APC (clone RPA-T4). Total cell numbers are not available, since part of the tonsil is always retained by the Pathology Department as fixed tissue for diagnostic purposes (table S2). For Cohort 2, usable flow cytometry was not available from all fresh specimens.

Anti-human antibodies for AIM assay are listed here, by company, Thermo Fisher Scientific: CD19 e780 (clone HIB19), CD14 e780 (clone 61D3), CD16 e780 (clone eBioCB16), OX40 FITC (clone Ber-ACT35), CD25 PE-Cyanine 7 (clone BC96), CD4 PerCP-eFluor710 (clone SK3); Biolegend: CD45RA BV570 (clone HI100), CXCR5 BV421 (clone J252D4), PD-1 BV785 (clone EH12.2H7), PD-L1 PE (clone 29E.2A3), CCR7 APC (clone G043H7) (table S3).

Anti-human antibodies for the proliferation assay using HLA class II cell lines are listed here, by company, Thermo Fisher Scientific: OX40 FITC (clone Ber-ACT35), CD25 PE-Cyanine 7 (clone BC96), CD4 PerCP-eFluor710 (clone SK3); Biolegend: PD-1 BV785 (clone EH12.2H7), PD-L1 PE (clone 29E.2A3) (table S4). Annexin V APC Apoptosis Detection kit was utilized (Thermofisher).

Anti-human antibodies for the granzyme B Assay are listed here, by company, Thermo Fisher Scientific: CD19 e780 (clone HIB19), CD14 e780 (clone 61D3), CD16 e780 (clone eBioCB16), OX40 PE (clone Ber-ACT35), CD25 PE-Cyanine 7 (clone BC96), CD4 PerCP-eFluor710 (clone SK3); Biolegend: CD45RA BV570 (clone HI100), CXCR5 BV421 (clone J252D4), PD-1 BV785 (clone EH12.2H7), Granzyme B Alexa Fluor 647 (clone GB11), and Alexa Fluor 647 Mouse IgG1, κ Isotype Control (clone MOPC-21) (table S5). Cells were acquired on a BD Fortessa and analyzed using FlowJo Software, version 9.9.4.

Anti-human antibodies for sorting GC Tfh and B cells are listed here, by company, Thermo Fisher Scientific: CD19 e780 (clone HIB19), CD14 e780 (clone 61D3), CD16 e780 (clone eBioCB16), CD8 e780 (clone RPA-T8), CD4 PerCP-eFluor710 (clone SK3), CD38 APC (clone HIT2); Biolegend: CD45RA BV570 (clone HI100), CXCR5 BV421 (clone J252D4), PD-1 BV785 (clone EH12.2H7), CCR7 BV650 (clone G043H7), CD20 BV570 (clone 2H7) (table S6). Anti-human antibodies for staining after a 5 day *in vitro* culture are listed here by company, Thermo Fisher Scientific: CD4 PerCP-eFluor710 (clone SK3), OX40 PE (clone Ber-ACT35), CD25 PE-Cyanine 7 (clone BC96), Biolegend: CD45RA BV570 (clone HI100), CXCR5 BV421 (clone J252D4), PD-1 BV785 (clone EH12.2H7), CD20 BV570 (clone 2H7), Granzyme B Alexa Fluor 647 (clone GB11), Perforin FITC (clone B-D48) (table S7). Cells were acquired on a BD Celesta and analyzed using FlowJo Software, version 9.9.4.

Anti-human antibodies for sorting for the cytotoxicity assay are listed here, by company, Thermo Fisher Scientific: CD14 e780 (clone 61D3), CD16 e780 (clone eBioCB16), CD4 PerCP-eFluor710 (clone SK3), CD8 PeCy7 (RPA-T8), PD-1 PE (clone eBioJ105), CD38 APC (clone HIT2), CD19 AF488 (HIB19); Biolegend: CCR7 BV650 (clone G043H7), CXCR5 APC (clone J252D4); BD Biosciences: CD45RA PE-CF594 (clone H100) (table S8). Cells were sorted on the BD FACSAria III or BD FACSAria Fusion. Data was analyzed using FlowJo 9.9.4.

### Histology

A small section was taken from each tonsil, fixed in 10% zinc formalin fixative for 24 hours at room temperature and transferred to 70% ethanol. For each tonsil, the microscopy core prepared a paraffin embedded section and an H&E stain. Slides were viewed using a Nikon Eclipse 80i. Images of three different locations on the same slide were taken (10X objective) and averaged per tonsil. The number of GCs and GC area were determined using “Image J” (NIH). Immunohistochemistry was performed by HistoTox Labs, Inc. (Boulder, Colorado). Each tissue was sectioned, mounted on a slide, and stained separately for CD20, Ki67, CD4, and PD-1.

### Immunofluorescence microscopy

A small section was taken from each tonsil, fixed in in 4% paraformaldehyde at 4°C for 2 hours, washed in PBS x 3 for 10 minutes, and placed in a 30% sucrose gradient for at least 18 hours at 4°C until the tissue sinks. The tissue section was washed in PBS and embedded in OCT compound using methylbutane and liquid nitrogen. Embedded tissues samples were stored at −80°C. Tissue sections were prepared by the LJI Microscopy core. For staining, slides were dried on the grill of the tissue culture hood for 30 minutes, washed in PBS x 2 for 10 minutes, and blocked with 10% FBS containing 0.5% Triton X-100 for 1 hour at room temperature. Antibodies were from Biolegend: CD4 Alexa Fluor 488 (clone RPA-T4) and Granzyme B Alexa Fluor 647 (clone GB11) and Isotype Control Alexa Fluor 647. Slides were stained overnight at 4°C. The next morning, slides were washed in PBS x 2 x 10 minutes, counterstained with Hoechst 3342 for 10 minutes, and washed in PBS x 2 x 10 minutes. Slides were then mounted in Prolonged Gold. Slides were visualized using Olympus FluoView FV10i Confocal.

### HLA typing

Genomic DNA was isolated from frozen tonsillar mononuclear cells using standard techniques (REPLI-g, Qiagen). Typing was performed at Murdoch University (Perth, Western Australia) *(53)*. We performed typing on the entire cohort; however, a few samples did not have enough DNA amplification to yield HLA results.

### Superantigen binding assay

Recombinant SpeA produced in *E. coli* (Toxin Technology) was biotinylated following manufacturer’s protocol using an EZ-Link Sulfo-NHS Biotinylation kit (Thermofisher). Biotinylated recombinant SpeA was incubated for 30 minutes at 4°C in FACS buffer using cell lines expressing different HLA receptors*(54, 55)*. DQB1*03:02 and DQB1*06:02 were expressed on the RM3 line. Cells were washed twice in FACS buffer. Streptavidin Alexa Fluor 647 (Biolegend) was used as a secondary stain. Cells were also labeled with fixable viability dye eFluor 780 (Thermo Fisher Scientific). Cells were fixed in 2% paraformaldehyde and acquired on BD FACS LSRII. Data was analyzed using FlowJo 9.9.4 and histograms generated using FlowJo 10.2.

### Superantigen stimulation assay

Antigen Presenting Cells (APCs): HLA Class II cell lines were cultured in R10 media containing RPMI, Penicillin/Streptomycin, L-Glutamax, 10% FBS, MEM Non-essential amino acids, and Sodium Pyruvate. For HLA DRB1 expressing L cell lines, the selection media included 200μg/mL G418. Prior to use of the HLA DRB1 cell line, 100μg/mL Butyric acid was added overnight to induce expression of the HLA DRB1 receptor. For HLA DQB1 expressing RM3 cell lines, the selection media included 12 μg/mL Blasticidin + 700 μg/mL G418. The number of APCs was optimized, using 5,000 cells per well of DRB1 expressing cell lines and 25,000 cells per well of DQB1 expressing cell lines. APCs were irradiated in a 96 well flat bottom tissue culture plate. CD4^+^ T cells: Cryopreserved PBMCs containing the HLA receptor of interest were thawed and purified using the EasySep Human CD4^+^ T cell enrichment kit, according to manufacturer’s protocol to 95 to 98% purity). CD4^+^ T cells were Cell Trace Violet (CTV) labeled and cultured at 100,000 cells per well. rSpeA was added to the well at different concentrations. As a control, CD4^+^ T cells alone were incubated with rSpeA in media consisting of RPMI + 10% Human AB sera (off the clot, Gemini) + penicillin/streptomycin + L-Glutamax. After 5 days, cells were analyzed for upregulation of activation marker OX40 and CTV.

### Antigen-specific CD4^+^ T cell assays

Tonsillar mononuclear cells were cultured at 1×10^6^ cells/well in AIM-V media in a 96 well round bottom plate for 18 hours. For the GAS-specific CD4^+^ T cell assay, cells were left unstimulated or stimulated with 10 μg/mL heat-inactivated, antibiotic-killed GAS*(25)*. As a comparison, cells were also stimulated with 10 μg/mL antibiotic-killed GAS or 10 μg/mL antibiotic-killed GAS deficient in SpeA, all from the same strain. The Nizet laboratory provided GAS strain M1T1 5448, originally isolated from a patient with necrotizing fasciitis and toxic shock syndrome*(56)*. A nonpathogenic Streptococcaceae, *Lactococcus lactis* NZ9000*(57)*, was used as a negative control. For the AIM assay, tonsillar cells were stimulated with 10 μg/mL antibiotic-killed *Lactococcus.* Bacteria were cultured in 100 mL Todd-Hewitt broth (Difco) statically at 37°C to OD600 0.6. Tissue culture grade penicillin/streptomycin (Invitrogen) was added to 1% and incubated for 1 hour. Cells were pelleted by centrifugation for 10 min at 4000xg, washed once and suspended in PBS. Samples were plated on Todd-Hewitt agar to confirm bacterial killing*(58)*. Total protein was quantified by bicinchonic acid assay (Pierce) for use as antigen. To inactivate superantigen, antibiotic-killed GAS was heat-treated at 65°C for 20 min. For the SpeA AIM assay, we utilized 1 μg/mL of rSpeA as a stimulus.

### Intracellular cytokine staining for granzyme B expression

Tonsillar mononuclear cells were cultured at 1×10^6^ cells/well in AIM-V media in a 96 well round bottom plate for 24 hours. Cells were either left unstimulated or stimulated with 1 μg/mL SpeA (Toxin Technology). At 20 hours, GolgiPlug was added prior to harvesting the cells at 24 hours for analysis, according to manufacturer’s protocol (BD Biosciences). Cells were permeabilized using the BD Cytofix/Cytoperm kit for intracellular cytokines.

### Intranuclear staining

Tonsillar mononuclear cells were cultured at 1×10^6^ cells/well in AIM-V media in a 96 well round bottom plate for 24 hours. Cells were permeabilized using the eBioscience Transcription buffer staining set (Thermofisher). FoxP3 PE (clone 236A/E7, Thermofisher) and Helios PE Dazzle (clone 22F6, Biolegend) were used.

### RNA sequencing

Tonsillar mononuclear cells were cultured at 1×10^6^ cells/well in AIM-V media in a 96 well round bottom plate for 18hours. Cells were stained using antibodies listed in Supplementary Table 2 with the exception of CCR7 and PD-L1. Cells were sorted on the BD FACSAria III or BD FACSAria Fusion for CD25^+^OX40^+^ GC Tfh cells. From 10 donors, cell numbers obtained ranged from 10^4^ to 10^5^ cells.

As described previously, total RNA was purified using a miRNAeasy micro kit (Qiagen) and quantified, as described previously*(59)*. Standard quality control steps were included to determine total RNA quality using Agilent Bioanalyzer (RNA integrity number (RIN) > 8.5; Agilent RNA 6000 Pico Kit)*(26, 60)*. Purified total RNA (0.25 to 5 ng) was amplified following the Smart-Seq2 protocol*(61)*. cDNA was purified using AMPure XP beads (1:1 ratio; Beckman Coulter). From this step, 1 ng cDNA was used to prepare a standard Nextera XT sequencing library (Nextera XT DNA sample preparation kit and index kit; Illumina). Samples were sequenced using a HiSeq2500 (Illumina) to obtain 50-bp single-end reads. Both whole-transcriptome amplification and sequencing library preparations were performed in a 96-well format to reduce assay-to-assay variability. Quality control steps were included to determine total RNA quality and quantity, the optimal number of PCR preamplification cycles (15 cycles), and fragment size selection. Samples that failed quality control were eliminated from further downstream steps. Barcoded Illumina sequencing libraries (Nextera; Illumina) were generated utilizing the automated platform (Biomek FXp). Libraries were sequenced on the HiSeq2500 Illumina platform to obtain 50-bp single-end reads (TruSeq Rapid Kit; Illumina), generating a median of ~13.6 million mapped 50 bp reads per sample).

### RNA-Seq analysis

The single-end reads that passed Illumina filters were filtered for reads aligning to tRNA, rRNA, adapter sequences, and spike-in controls. The reads were then aligned to UCSC hg19 reference genome using TopHat (v 1.4.1)*(62)*. DUST scores were calculated with PRINSEQ Lite (v 0.20.3)*(63)* and low-complexity reads (DUST > 4) were removed from the BAM files. The alignment results were parsed via the SAMtools*(64)* to generate SAM files. Read counts to each genomic feature were obtained with the htseq-count program (v 0.6.0)*(65)* using the “union” option. After removing absent features (zero counts in all samples), the raw counts were converted to RPKM values and filtered by setting a cutoff value of 1. Multiplot Studio in the GenePattern suite (http://www.broadinstitute.org/cancer/software/genepattern/) was employed to generate the volcano plot with RPKM values. The raw counts were then imported to R/Bioconductor package DESeq2*(66)* to identify differentially expressed genes among conditions. DESeq2 normalizes counts by dividing each column of the count table (samples) by the size factor of this column. The size factor is calculated by dividing the samples by geometric means of the genes. This brings the count values to a common scale suitable for comparison. P-values for differential expression are calculated using Wald test that estimates the significance of coefficients in a fitted negative binomial generalized linear model (GLM). These p-values are then adjusted for multiple test correction using Benjamini Hochberg algorithm*(67)* to control the false discovery rate. Cluster analyses including principal component analysis (PCA) and hierarchical clustering were performed using standard algorithms and metrics. Hierarchical clustering was performed using complete linkage with Euclidean metric.

### Sorting GC Tfh and non-GC B cells for Granzyme B expression

Tonsillar mononuclear cells were sorted (antibodies listed in Supplementary Table 6) for GC Tfh (CXCR5^hi^PD-1^hi^ of CD45RA^-^CD4^+^) and non-GC B cells (CD20^+^CD38^-^) to serve as APCs. Cells were plated at 75,000 GC Tfh and GC B cells 96 well round bottom plates in media containing 10% human sera (RPMI + penicillin/streptomycin + L-glutamax) + IL-7 (Final concentration 4ng/mL). Cells were either left unstimulated or stimulated with 1 μg/mL SpeA. After a 5 day *in vitro* culture, cells were harvested and stained for granzyme B (antibodies listed in Supplementary Table 7).

### Cytotoxicity assay

Tonsillar mononuclear cells were sorted (antibodies listed in Supplementary Table 8) for GC Tfh (CXCR5^hi^PD-1^hi^ of CD45RA^-^CD4^+^), mTfh (CXCR5^+^PD-1^+^ of CD45RA^-^CD4^+^), non-Tfh (CXCR5^-^ of CD45RA^-^CD4^+^), naïve CD4^+^ (CCR7^+^CD45RA^+^), and CD8^+^ T cells as effector cells. Tonsillar mononuclear cells were also sorted for autologous non GC B and plasma cells (CD19^+^CD38^-^) to serve as target cells. B cells were labeled with CTV and cultured at a 2:1 ratio of effector cells to 1 target cell in media containing 5% human sera (RPMI + penicillin/streptomycin + L-glutamax)*(68)*. Cells were plated at 50,000 target cells to 100,000 effector cells in 96 well round bottom plates. Cells were either left unstimulated or stimulated with 1 μg/mL SpeA. As a control, B cells were also left unstimulated or stimulated with 1 μg/mL SpeA. After 40 hours of incubation, cells were harvested and the number of CTV^+^ cells was quantified by flow cytometry*(69)*. Cells were plated at least in triplicate, depending on how many GC Tfh cells were sorted from each tonsil. Killing capacity for GC Tfh cells was determined by averaging the absolute counts of CTV-labeled B cells co-cultured with unstimulated GC Tfh or naïve CD4^+^ T cells = B. The absolute cell count of CTV-labeled B cells co-cultured with SpeA-stimulated effector cells was then determined for each well = A.

Equation: % Killing Capacity = [1 – (A/B)]*100

In some experiments, a blocking antibody to Fas Ab (EMD Millipore) and FasL Ab (R&D) were co-cultured during the cytotoxicity assay.

### ImageStream

Images were acquired on a 2-camera ImageStream MkII imaging flow cytometer (Amnis, Seattle) with 60X objective and Inspire software version 200.1. The cytometer passed all ASSIST performance checks prior to image acquisition. FITC (Ch02, 480-560 nm), PE (Ch03, 560-595 nm) PE-CF594 (Ch04, 595-642 nm), PerCP-eFluor710 (Ch05, 648-745 nm) and PE-Cy7 (Ch06, 745-780 nm) were excited at 488nm (200 mW). BV421 (Ch07, 435-505 nm) and BV510 (Ch08, 505-570 nm) were excited at 405 nm (120 mW). APC (Ch11, 640-745 nm) and APC-eFluor780 (Ch12, 745-780 nm) were excited at 642 nm (150 mW). 10,000 single, in-focus, dump-negative, CD3-positive events were acquired per sample. Data were compensated and analyzed with IDEAS software version 6.2 using the default masks and feature set.

### ELISA

Plasma from RT and non-RT children was tested for IgG, Streptolysin O (SLO) IgG and SpeA IgG. To determine IgG titer, human IgG antibody was coated (1:5000 dilution in PBS) overnight. To determine SLO IgG titer, recombinant Streptolysin O (Abcam) produced in *E. coli* was coated at 1 μg/mL. To determine SpeA IgG titer, recombinant SpeA (Toxin Technologies) was coated at 1 μg/mL. Plates were coated overnight at 4°C. PBS + 0.05% Tween was used for all washes. Plates were blocked with PBS containing 0.2% Tween and 1% BSA at room temperature for 90 minutes. For IgG, human IgG was utilized as a standard. For SpeA and SLO, pooled plasma from normal healthy human donors was utilized as a standard to establish “relative units” of SpeA and SLO IgG in RT and non-RT plasma. As a secondary, a monoclonal mouse anti-human IgG antibody conjugated to HRP (Hybridoma Reagents Laboratory) was used.

### Statistical analysis

All statistical analyses were performed using two-tailed Mann Whitney test using a nonparametric distribution in GraphPad 7.0, unless otherwise specified. Two-tailed Fisher exact test was determined using either GraphPad software or R software version 3.3.1.

## ACKNOWLEDGEMENTS

Supported by a Thrasher Research Fund for an Early Career Award (JD), the US National Institutes of Health (Center for HIV/AIDS Vaccine Immunology and Immunogen Discovery Grant 1UM1AI100663, S.C.), internal funding from the La Jolla Institute (SC), and the UCSD Department of Medicine Division of Infectious Diseases NIH Training Grants (JD, 5T32AI007036-35 and 5T32AI007384-25), NIAID grant AI077780 (VN), Clinical Research Fellowship Grant UK Charity No. 1089464 (DL, RC, CO), NIH S10OD016262 (LJI), and NIH S10RR027366 (LJI).

We thank Scott D. Boyd, Katherine J.L. Jackson, and Jean Oak at Stanford University Medical Center for assistance with histology. We thank Cheryl Kim, Lara Nosworthy, Denise Hinz, and Robin Simmons at LJI for assistance with flow cytometry sorting. We thank Zbigniew Mikulski, Angela Lamberth, and the LJI Microscopy Core for assistance with confocal imaging. We thank Yoav Altman, the Sanford Burnham Prebys Flow Cytometry Shared Resource, and the James B. Pendleton Charitable Trust for assistance with Image Stream. We thank Amy Baxter at the University of Montreal Hospital Research Center for further assistance with Image Stream software. We thank Ashish Mittal at UCSD School of Medicine for help with gathering clinical data and Nour Bundogji at Rady Children’s Hospital for help with study enrollment. We thank the RNA sequencing core and Alex Fu at LJI for assistance with bioinformatics. We thank Michela Locci with assistance with bioinformatics analysis. We thank Carla Oseroff for help with the HLA cell lines. We thank Daniela Weiskopf at LJI and Daniel Kaufmann at the University of Montreal Hospital Research Center with advice for CD4^+^ T cell cytotoxicity assay. We thank April Frazier and John Sidney with assistance with HLA typing and analyses. We thank the donors for their participation.

## AUTHOR CONTRIBUTIONS

S.C., C.H., and J.M.D. designed the study. M.B. and M.B. enrolled patients and provided clinical specimens. K.Ke. and K.Ka. assisted with tonsillar cell isolation. K.Ke. measured GC areas and performed immunohistochemistry. J.M.D. analyzed immunophenotyping and HLA-typing data. C.L.A. enrolled patients and provided HLA-typed PBMCs. A.S. helped facilitated HLA-typing and provided HLA expressing cell lines. E.A., C.L., V.N. provided antibiotic-killed GAS, antibiotic-killed *Lactococcus lactis*, and SpeA-deficient GAS. J.M.D. designed the AIM assay, performed flow cytometry and confocal microscopy experiments. S.R., P.V., and G.S. performed RNA-sequencing. D.L., R.C., and C.O. enrolled patients for lymph node studies and provided healthy lymph nodes. S.C. and J.M.D. wrote the manuscript with input from all authors.

## Competing Financial Interests

The authors declare no competing financial interests.

---

**Supplementary Figure 1.**
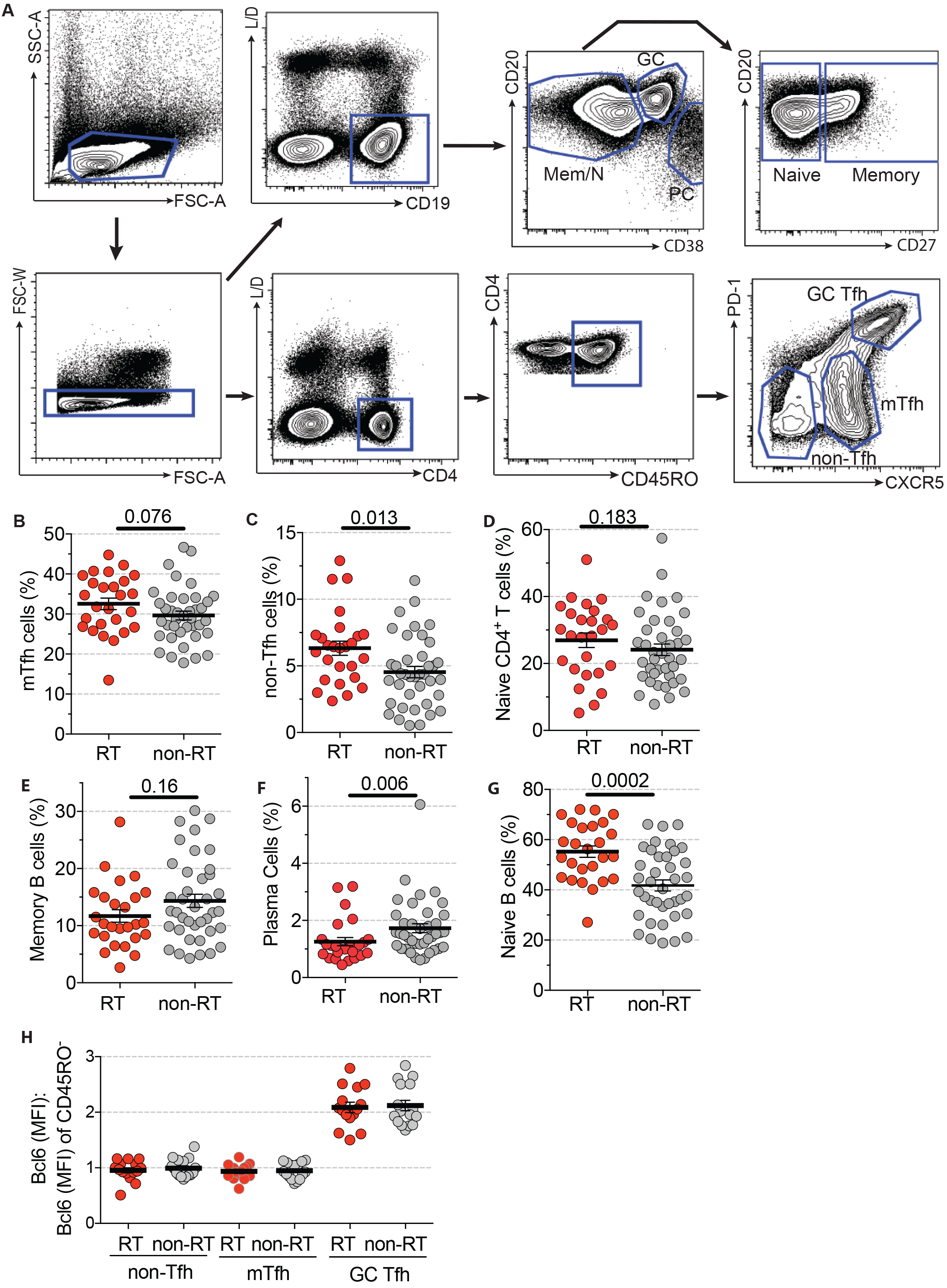
Immunophenotyping RT and non-RT tonsils. **(A)** Gating strategy for tonsillar CD4^+^ T cells and B cells. **(B)** RT tonsils (n=26) have significantly more mTfh CD4^+^ T cells (CXCR5^+^PD-1^+^ of CD45RO^+^CD4^+^) and **(C)** non-Tfh CD4^+^ T cells (CXCR5^−^ of CD45RO^+^CD4^+^) than non-RT tonsils (n=39). mTfh and non-Tfh cells were gated on antigen-experienced (CD45RA^−^CD4^+^) T cells and normalized to CD4^+^ T cells. **(D)** There is no difference in frequency of naive (CD45RO^−^) CD4^+^ T cells. **(E)** There is no difference in memory B cells (CD27^+^CD20^+^ of CD19^+^). **(F)** RT tonsils have significantly fewer plasma cells (CD38^hi^CD20^hi^ of CD19^+^) than non-RT tonsils. **(G)** There are significantly more naive B cells (CD27^−^CD20^+^ of CD19^+^) in RT tonsils than non-RT tonsils. Statistical significance determined by Mann Whitney test. **(H)** There is no difference in BCL6 expression by GC Tfh cells from RT (n=15) and non-RT tonsils (n=16), *P*=0.98. BCL6 MFI was quantified for GC Tfh, mTfh, non-Tfh, and CD45RO^−^(naïve) CD4^+^ T cells. The MFI of BCL6 for GC Tfh, mTfh, and non-Tfh was then normalized to the MFI of BCL6 in CD45RO^−^ CD4^+^ T cells.
Statistical significance determined by Mann-Whitney test (b-h).

**Supplementary Figure 2.**
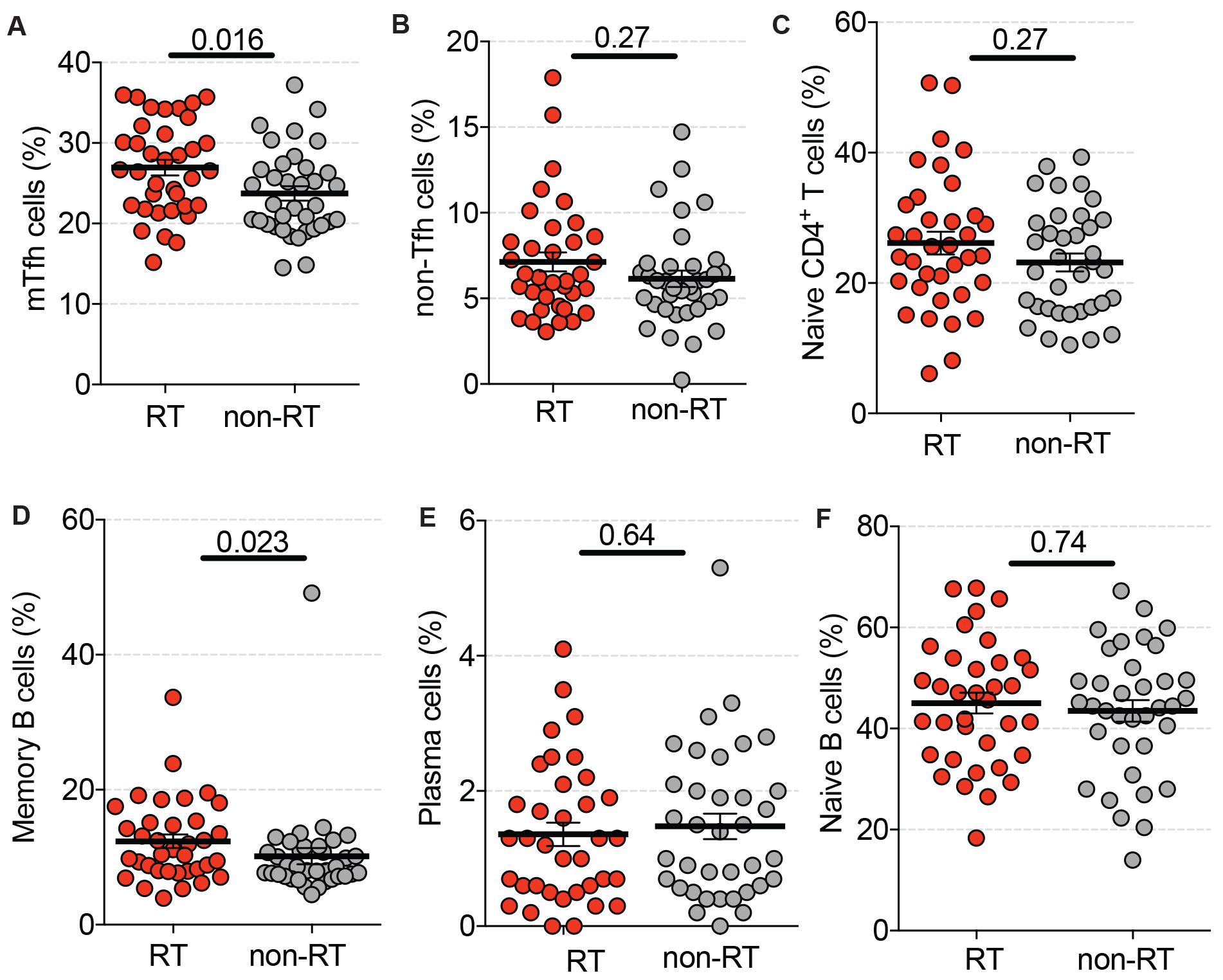
Immunophenotype of tonsils from the 2^nd^ Cohort. **(A)** RT tonsils (n=36) have significantly more mTfh CD4^+^ T cells (CXCR5^+^PD-1^+^ of CD45RO^+^CD4^+^) and **(C)** non-Tfh CD4^+^ T cells (CXCR5^−^ of CD45RO^+^CD4^+^) than non-RT tonsils (n=36). mTfh and non-Tfh cells were gated on antigen-experienced (CD45RA^−^CD4^+^) T cells and normalized to CD4^+^ T cells. **(D)** There is no difference in memory B cells (CD27^+^CD20^+^ of CD19^+^). **(E)** RT tonsils have significantly fewer plasma cells (CD38^hi^CD20^hi^ of CD19^+^) than non-RT tonsils. **(F)** There are significantly more naive B cells (CD27^−^CD20^+^ of CD19^+^) in RT tonsils than non-RT tonsils. Statistical significance determined by Mann Whitney test.

**Supplementary Figure 3.**
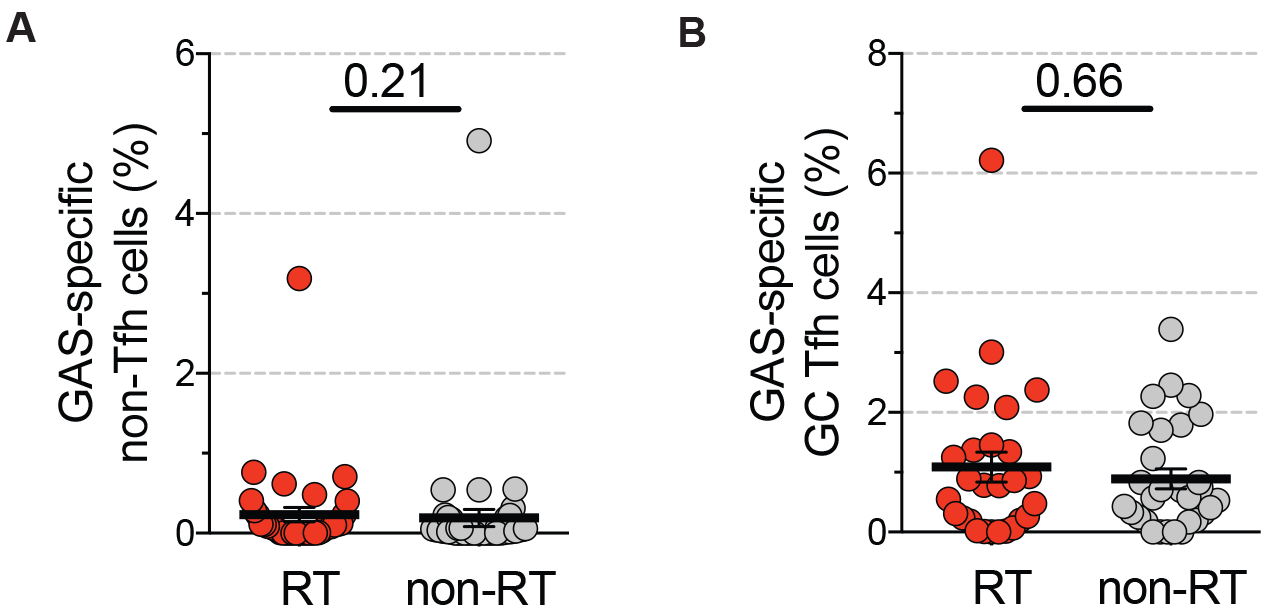
GAS-specific CD4^+^ T cells in RT and non-RT tonsils. **(A)** RT tonsils and non-RT tonsils contained comparable frequencies of GAS-specific non-Tfh cells. GAS-specific non-Tfh cells are quantified as % of total CD4^+^ T cells. **(B)** RT tonsils and non-RT tonsils contained comparable frequencies of GAS-specific GC Tfh cells. GAS-specific GC Tfh cells are quantified as % of total CD4^+^ T cells. Statistical significance determined by Mann-Whitney test (a,b).

**Supplementary Figure 4.**
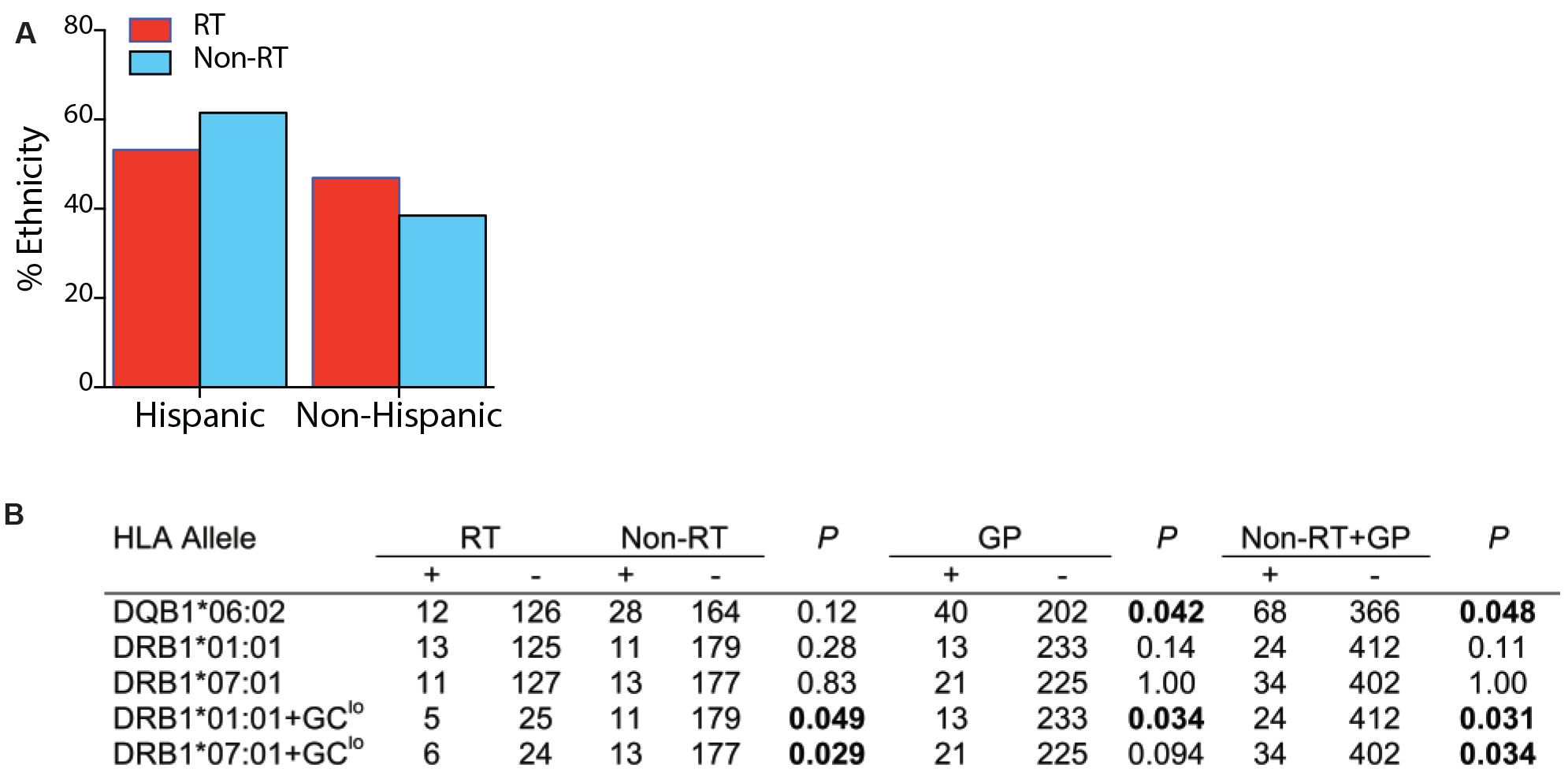
RT and non-RT patient HLA types. **(A)** Percentage of Hispanic and non-Hispanics among children with RT and non-RT. **(B)** Allelic frequencies in RT, non-RT, GP, and GP+non-RT individuals for HLA class II alleles of interest. P values represent comparison between RT and non-RT, RT and GP, and RT and GP+non-RT. Statistical significance determined by Fisher Exact Test.

**Supplementary Figure 5.**
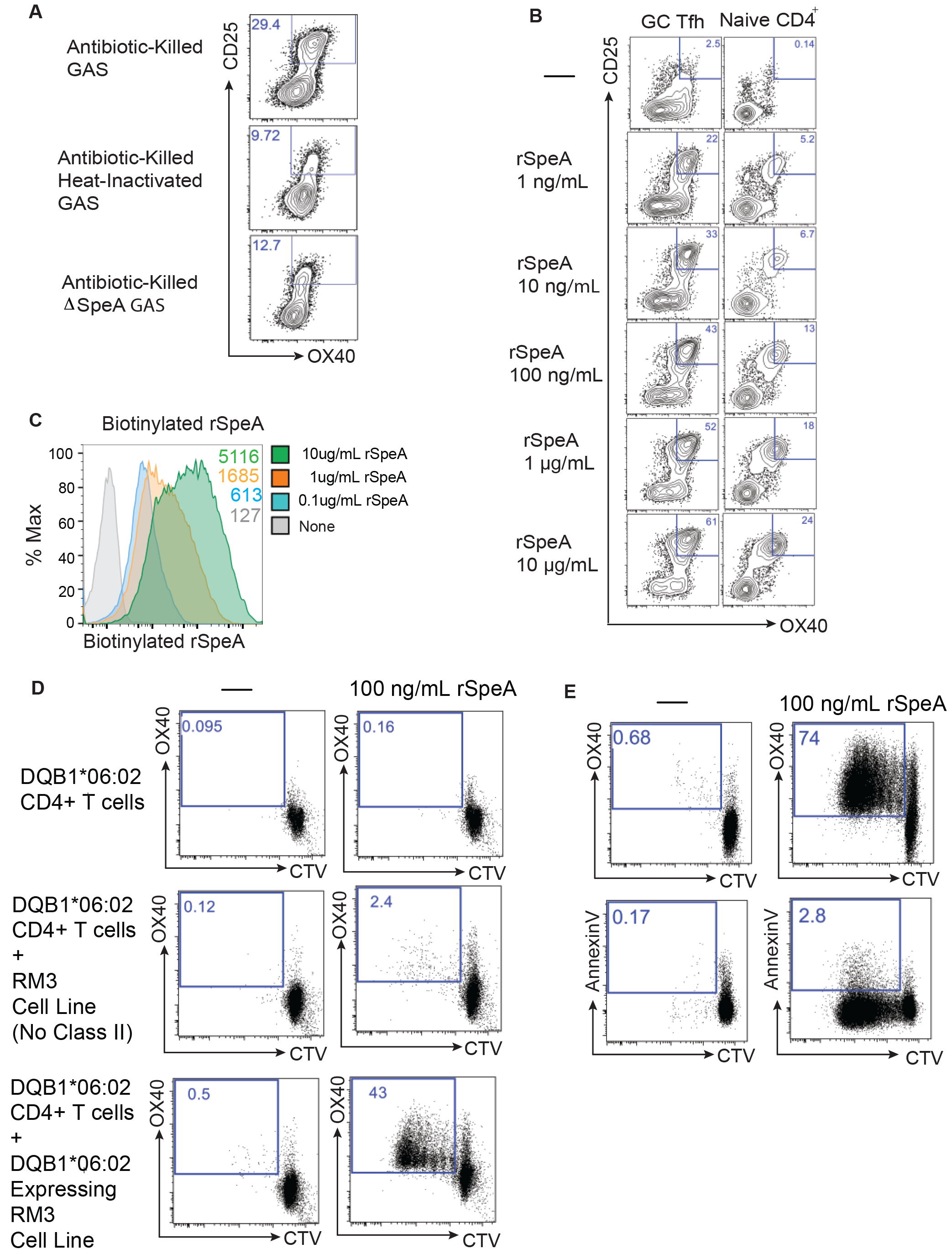
SpeA-responsive GC Tfh cells. **(A)** Flow cytometry gating of activated (AIM^+^) GC Tfh cells following stimulation with antibiotic-killed GAS, heat-inactivated antibiotic-killed GAS and antibiotic-killed GAS lacking SpeA. **(B)** Dose response of GC Tfh cells and naive CD4^+^ T cells to SpeA. **(C)** Dose-dependent binding of biotinylated SpeA (recombinant SpeA, rSpeA) to an RM3 cell line expressing HLA DQB1*06:02. **(D)** Flow cytometry plots of unstimulated and SpeA-stimulated HLA DQB1*06:02 CD4^+^ T cells alone (top panel), HLA DQB1*06:02 CD4^+^ T cells co-cultured with the RM3 cell line, and HLA DQB1*06:02 CD4^+^ T cells co-cultured with the RM3 cell line expressing DQB1*06:02. **(E)** Flow cytometry plots of unstimulated and SpeA-stimulated DQB1*06:02 CD4^+^ T cells co-cultured with an RM3 cell line expressing HLA DQB1*06:02 and subsequently stained with Annexin V. There is minimal Annexin V expression by SpeA-responsive CD4^+^ T cells.

**Supplementary Figure 6.**
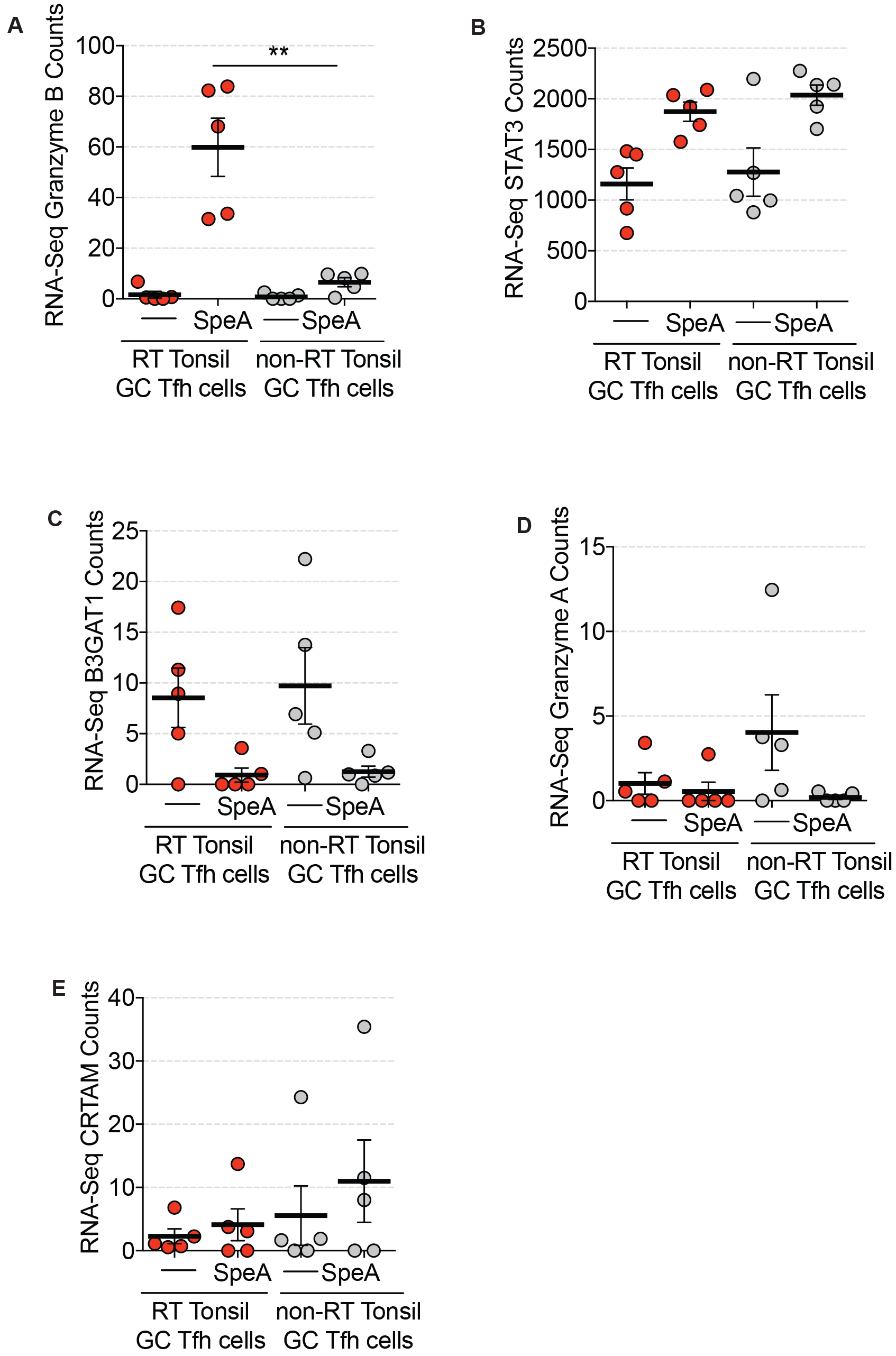
SpeA-responsive GC Tfh cells. Comparison of RNA-seq counts by CD4^+^ T cells from normal lymph nodes and blood, unstimulated GC Tfh cells, GAS-specific GC Tfh cells (AIM^+^), and SpeA-responsive GC Tfh cells (AIM^+^) by RT tonsils and non-RT tonsils. **(A)** Comparison of granzyme B counts. SpeA-responsive GC Tfh cells from RT tonsils expressed significantly more granzyme B RNA than non-RT tonsils. **(B)** Comparison of STAT3 counts. There is no difference between RT and non-RT tonsils. **(C)** Comparison of B3GAT1 counts. There is no difference between RT and non-RT tonsils. **(D)** Comparison of granzyme A counts. There is no difference between RT and non-RT tonsils. **(E)** Comparison of CRTAM counts. There is no difference between RT and non-RT tonsils. Statistical significance determined by Mann-Whitney test.

**Supplementary Figure 7.**
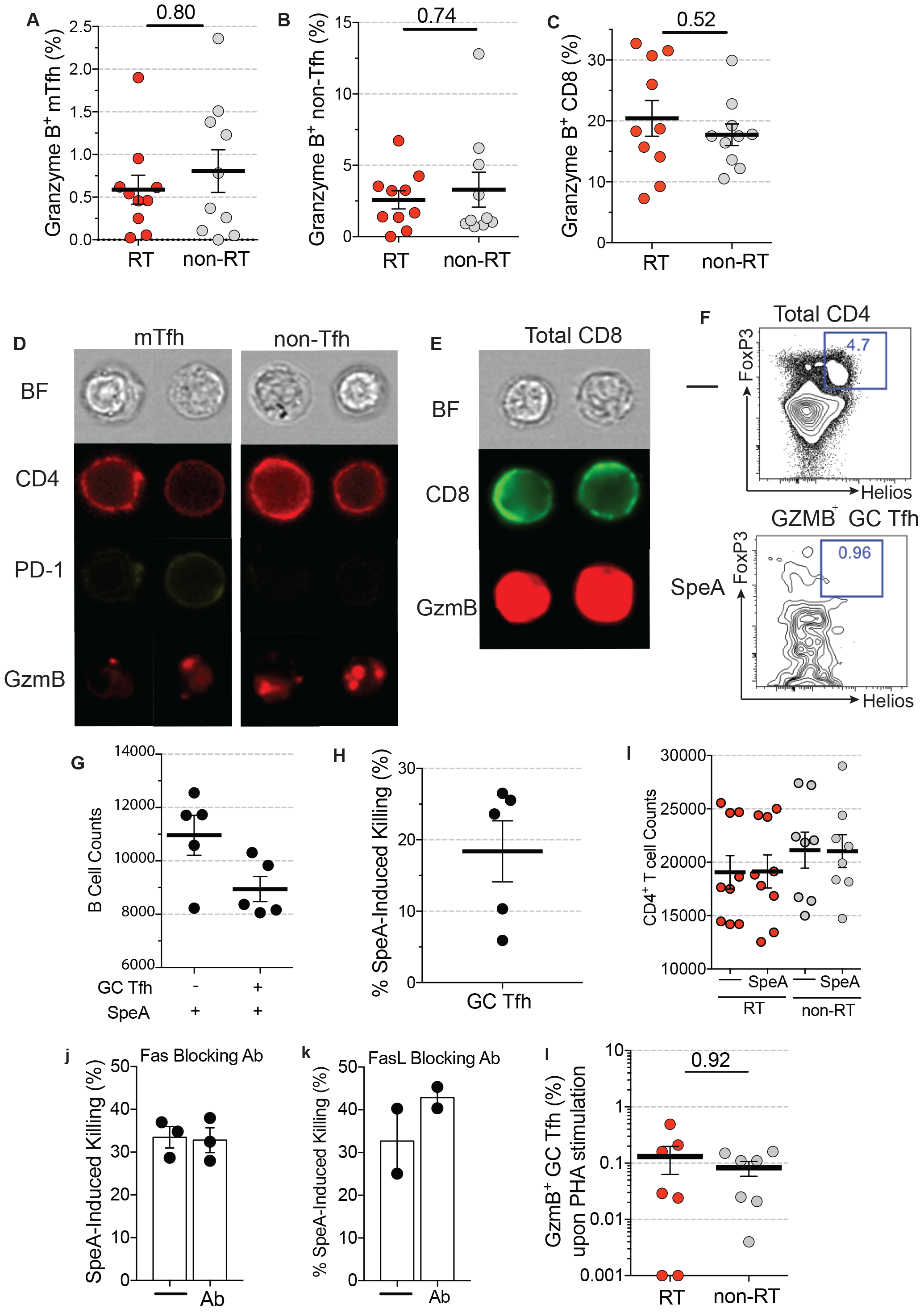
SpeA induced granzyme B production. Granzyme B expression by
**(A)** mTfh cells, **(B)** non-Tfh cells, and **(C)** CD8^+^ T cells from RT tonsils and non-RT tonsils following SpeA stimulation, as measured by flow cytometry. **(D)** ImageStream examples of granzyme B expression by SpeA-responsive mTfh cells, non-Tfh cells, and **(E)** CD8^+^ T cells from an RT tonsil. **(F)** Percentage of T follicular regulatory (Tfr) cells (FoxP3^+^Helios^+^) from unstimulated total CD4^+^ T cells, SpeA-stimulated total CD4^+^ T cells, and granzyme B^+^ GC Tfh cells. **(G)** B cell counts following 40h co-culture with GC Tfh cells, unstimulated or stimulated with SpeA. B cell death was not observed in the absence of SpeA stimulation. A representative donor is shown. **(H)** SpeA-induced cytotoxicity by GC Tfh (CXCR5^hi^PD-1^hi^CD45RA^−^CD4^+^) from the same donor as in **(G)** of autologous non-GC B cells (CD19^+^CD38^−^). **(I)** Cell counts of remaining GC Tfh cells following co-culture with B cells, left unstimulated or stimulated with SpeA. GC Tfh cells from 8 RT and 8 non-RT are shown. GC Tfh cell fratricide was not observed. **(J)** Monoclonal antibodies blocking Fas or **(K)** FasL did not inhibit the SpeA-induced GC Tfh cell killing of B cells. **(L)** Minimal granzyme B^+^ GC Tfh cells observed following stimulation with PHA. RT and non-RT. Statistical significance determined by Mann-Whitney test.

**Supplementary Table 1.**
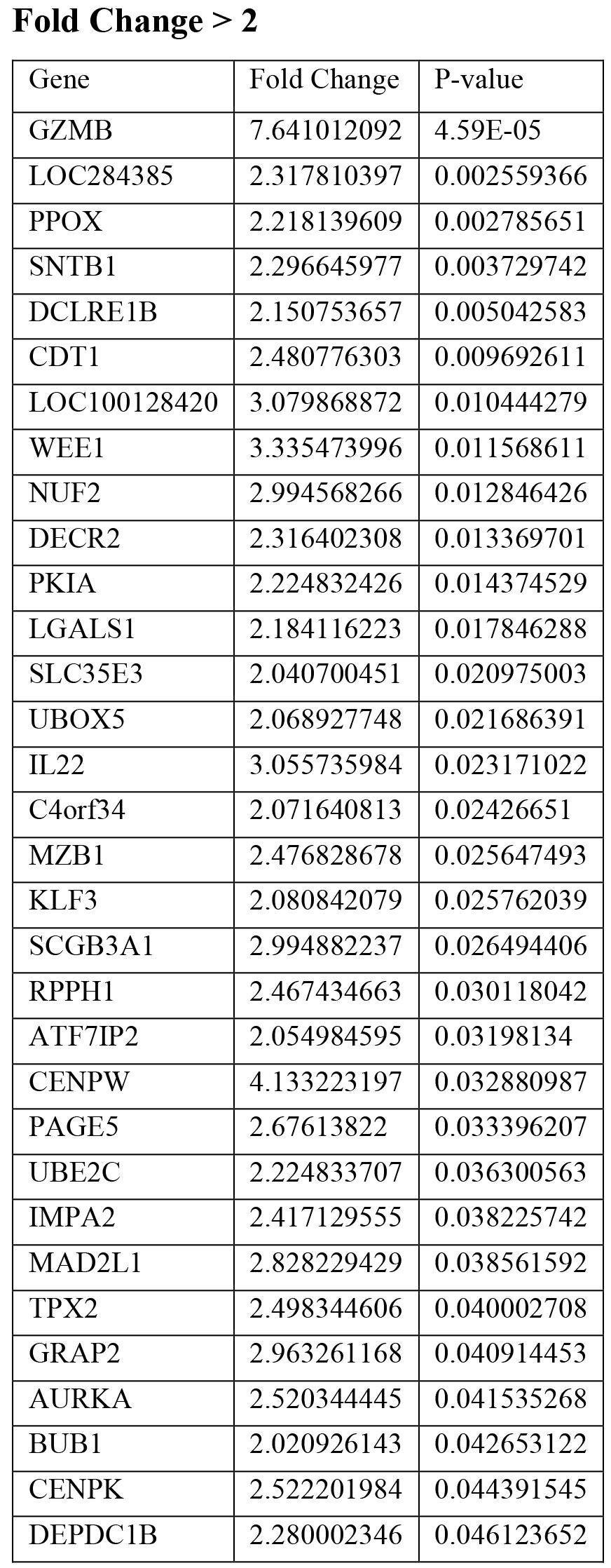
RNA-seq analysis of gene expression by SpeA stimulated GC Tfh cells from RT tonsils compared to non-RT tonsils, presented as reads per kilobase of transcript per million mapped reads (RPKM). Gene expression by SpeA stimulated GC Tfh cells is plotted against *P* values (RT over non-RT tonsils) relative to fold change in (RT over non-RT tonsils).

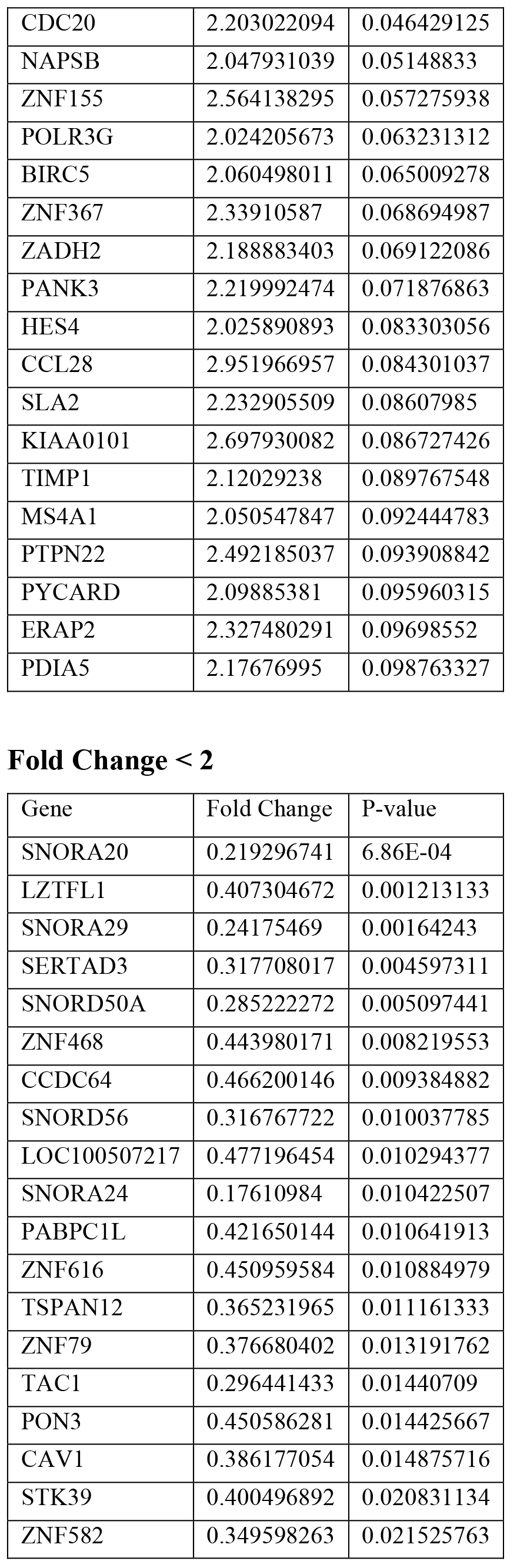

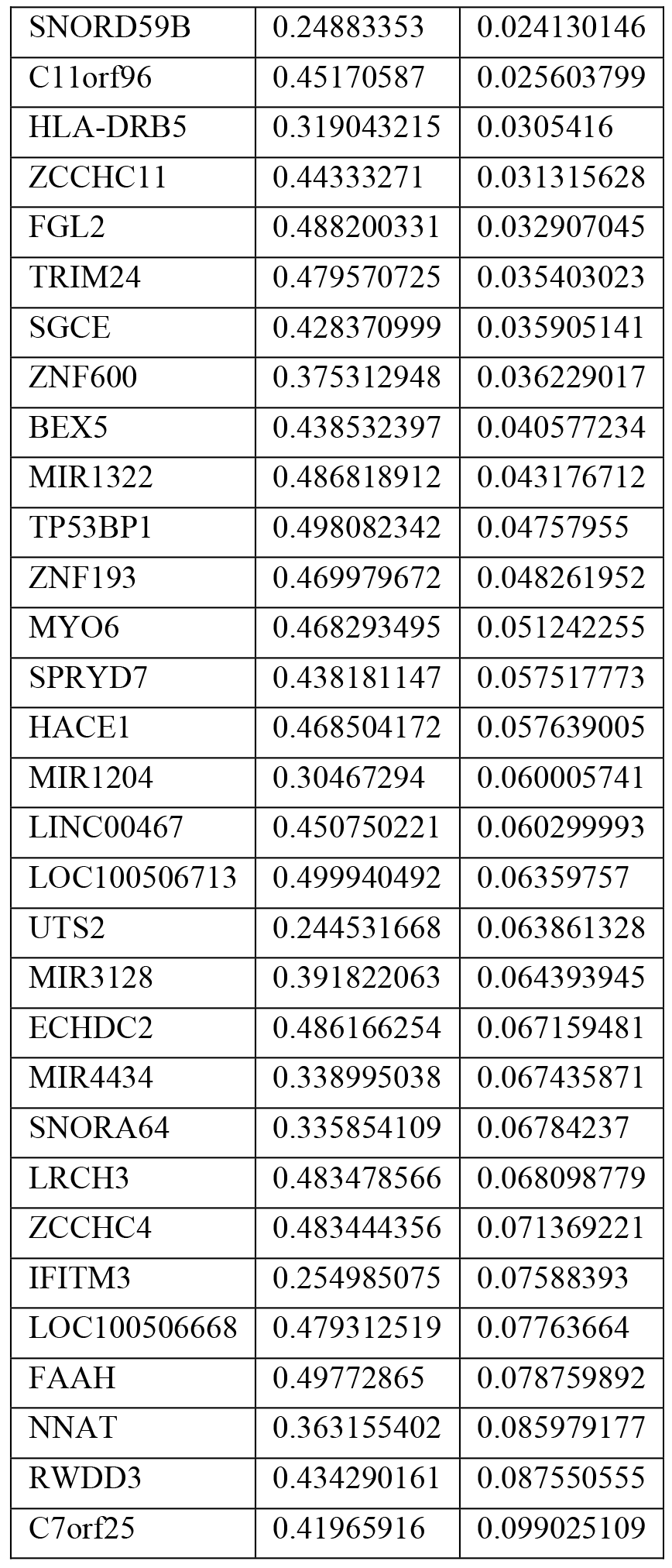

**Supplementary Table 2.**
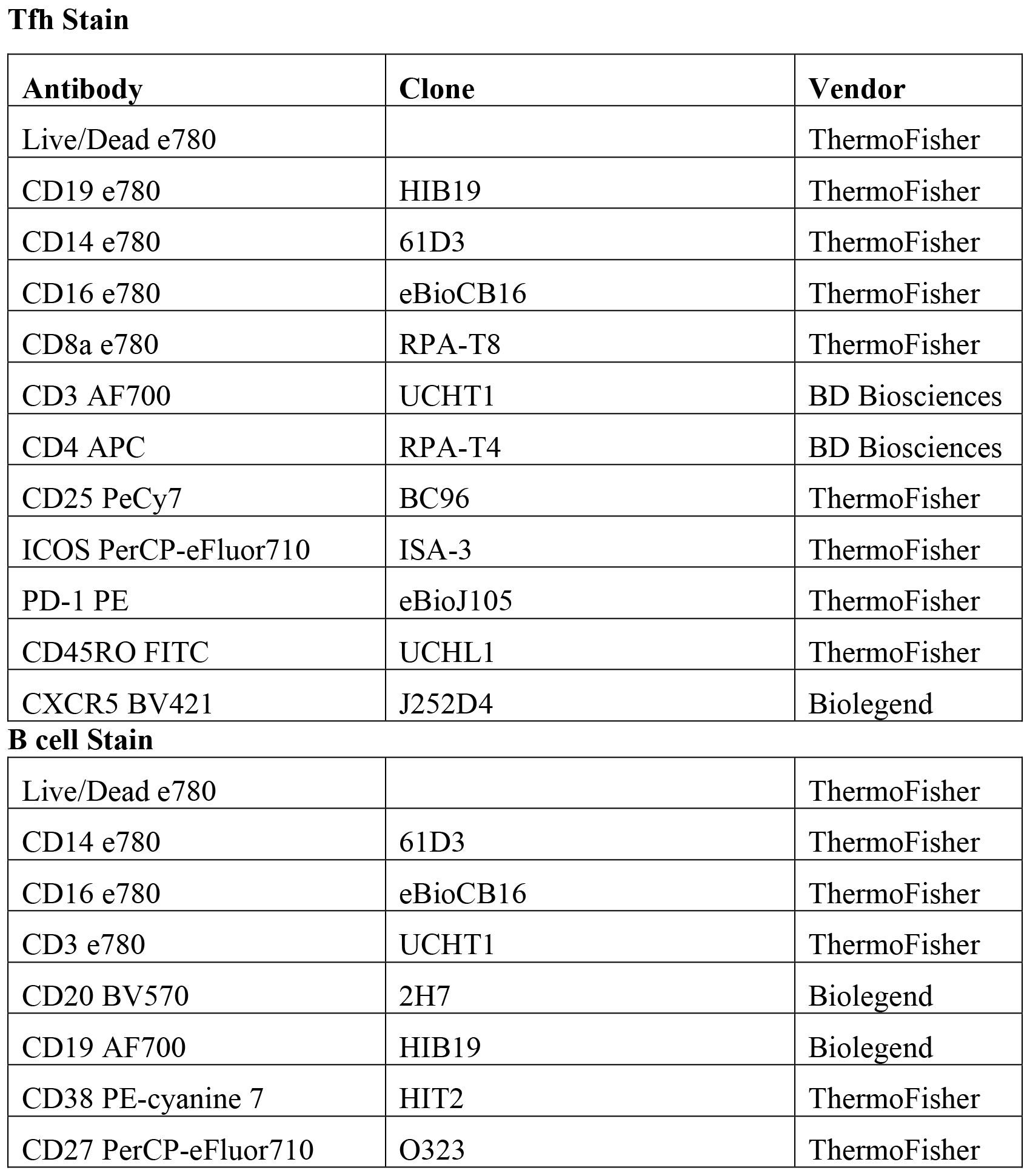
Flow cytometry antibodies for fresh tonsil stain

**Tfh Stain**
| **Antibody** | **Clone** | **Vendor** |
| --- | --- | --- |
| Live/Dead e780 |  | ThermoFisher |
| CD19 e780 | HIB19 | ThermoFisher |
| CD14 e780 | 61D3 | ThermoFisher |
| CD16 e780 | eBioCB16 | ThermoFisher |
| CD8a e780 | RPA-T8 | ThermoFisher |
| CD3 AF700 | UCHT1 | BD Biosciences |
| CD4 APC | RPA-T4 | BD Biosciences |
| CD25 PeCy7 | BC96 | ThermoFisher |
| ICOS PerCP-eFluor710 | ISA-3 | ThermoFisher |
| PD-1 PE | eBioJ105 | ThermoFisher |
| CD45RO FITC | UCHL1 | ThermoFisher |
| CXCR5 BV421 | J252D4 | Biolegend |
| **B cell Stain** |  |  |
| Live/Dead e780 |  | ThermoFisher |
| CD14 e780 | 61D3 | ThermoFisher |
| CD16 e780 | eBioCB16 | ThermoFisher |
| CD3 e780 | UCHT1 | ThermoFisher |
| CD20 BV570 | 2H7 | Biolegend |
| CD19 AF700 | HIB19 | Biolegend |
| CD38 PE-cyanine 7 | HIT2 | ThermoFisher |
| CD27 PerCP-eFluor710 | O323 | ThermoFisher |

**Supplementary Table 3.** Flow cytometry antibodies for AIM assay

| **Antibody** | **Clone** | **Vendor** |
| --- | --- | --- |
| Live/Dead e780 |  | ThermoFisher |
| CD19 e780 | H1B19 | ThermoFisher |
| CD14 e780 | 61D3 | ThermoFisher |
| CD16 e780 | eBioCB16 | ThermoFisher |
| CD8a e780 | RPA-T8 | ThermoFisher |
| CD45RA BV570 | HI100 | Biolegend |
| CCR7 APC | G043H7 | Biolegend |
| CD25 PeCy7 | BC96 | ThermoFisher |
| OX40 FITC | BER-ACT35 | Biolegend |
| PD-L1 PE | 29E.2A3 | Biolegend |
| CXCR5 BV421 | J252D4 | Biolegend |
| PD-1 BV785 | EH12.2H7 | Biolegend |
| CD4 PerCP-eFluor710 | SK3 | ThermoFisher |

**Supplementary Table 4.** Flow cytometry antibodies for PBMC Proliferation Assay

| **Antibody** | **Clone** | **Vendor** |
| --- | --- | --- |
| Live/Dead e780 |  | eBioscience |
| CD25 PeCy7 | BC96 | eBioscience |
| OX40 FITC | BER-ACT35 | Biolegend |
| PD-L1 PE | 29E.2A3 | Biolegend |
| PD-1 BV785 | EH12.2H7 | Biolegend |
| CD4 PerCP-eFluor710 | SK3 | ThermoFisher |

**Supplementary Table 5.** Flow cytometry antibodies for Granzyme B Detection

| **Antibody** | **Clone** | **Vendor** |
| --- | --- | --- |
| Live/Dead e780 |  | ThermoFisher |
| CD19 e780 | HIB19 | ThermoFisher |
| CD14 e780 | 61D3 | ThermoFisher |
| CD16 e780 | eBioCB16 | ThermoFisher |
| CD3 AF700 | UCHT1 | BD Biosciences |
| CD45RA BV570 | HI100 | Biolegend |
| CXCR5 BV421 | J252D4 | Biolegend |
| PD-1 BV785 | EH12.2H7 | Biolegend |
| CD4 PerCP-eFluor710 | SK3 | ThermoFisher |
| CD25 PeCy7 | BC96 | ThermoFisher |
| OX40 PE | BER-ACT35 | Biolegend |
| CD8a BV650 | RPA-T8 | Biolegend |
| Granzyme B AF647 | GB11 | Biolegend |
| Mouse IgG1, $\kappa$ Isotype Control AF647 | MOPC-21 | Biolegend |

**Supplementary Table 6.**
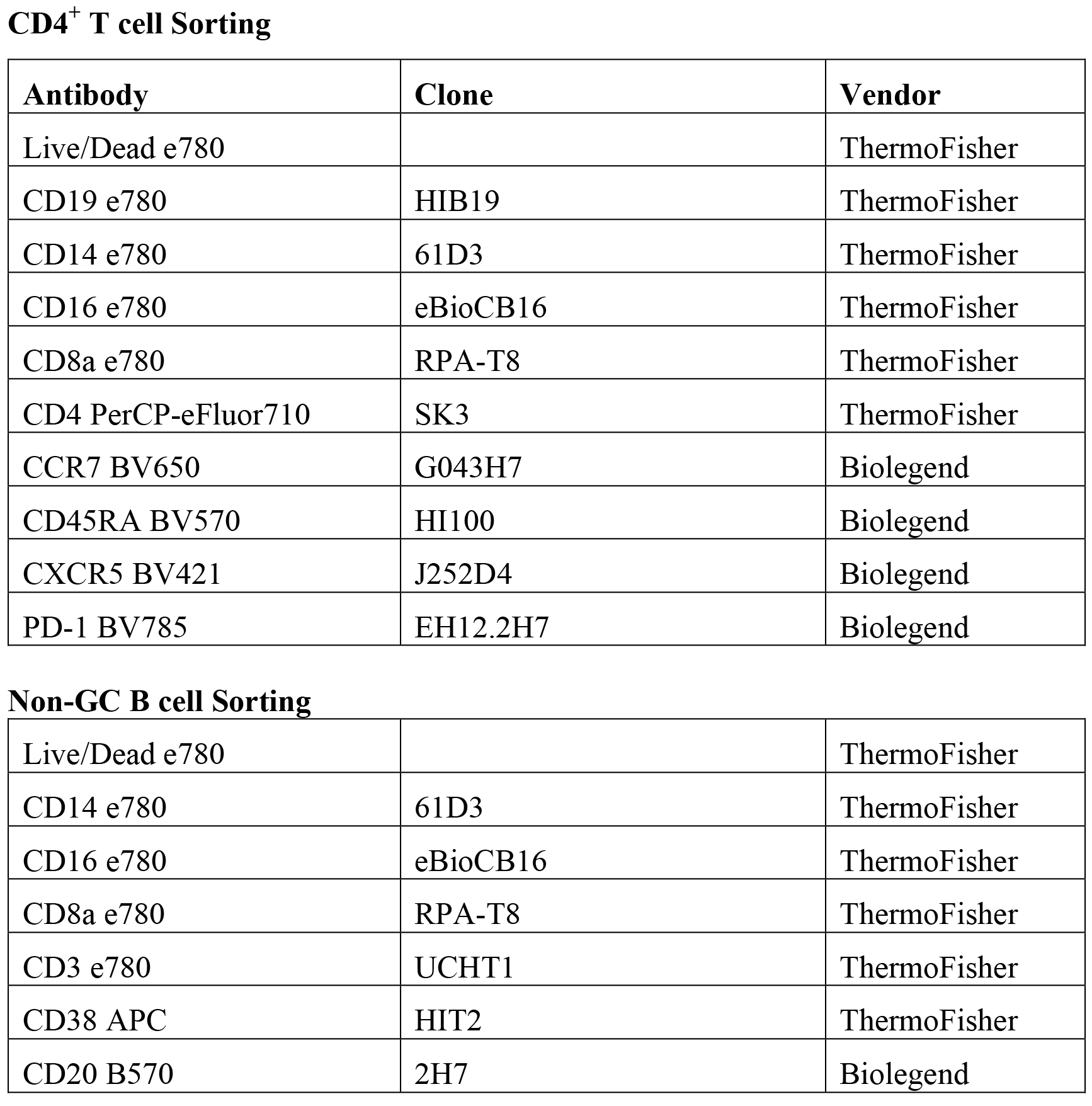
Flow cytometry antibodies for used for sorting GC Tfh and non-GC B cells for granzyme B expression after 5 day *in vitro* culture.

**CD4<sup>+</sup> T cell Sorting**
| Antibody | Clone | Vendor |
| --- | --- | --- |
| Live/Dead e780 |  | ThermoFisher |
| CD19 e780 | H1B19 | ThermoFisher |
| CD14 e780 | 61D3 | ThermoFisher |
| CD16 e780 | eBioCB16 | ThermoFisher |
| CD8a e780 | RPA-T8 | ThermoFisher |
| CD4 PerCP-eFluor710 | SK3 | ThermoFisher |
| CCR7 BV650 | G043H7 | Biolegend |
| CD45RA BV570 | HI100 | Biolegend |
| CXCR5 BV421 | J252D4 | Biolegend |
| PD-1 BV785 | EH12.2H7 | Biolegend |
| **Non-GC B cell Sorting** |  |  |
| Live/Dead e780 |  | ThermoFisher |
| CD14 e780 | 61D3 | ThermoFisher |
| CD16 e780 | eBioCB16 | ThermoFisher |
| CD8a e780 | RPA-T8 | ThermoFisher |
| CD3 e780 | UCHT1 | ThermoFisher |
| CD38 APC | HIT2 | ThermoFisher |
| CD20 B570 | 2H7 | Biolegend |

**Supplementary Table 7.** Flow cytometry antibodies for Granzyme B Detection from sorted GC Tfh cells.

| **Antibody** | **Clone** | **Vendor** |
| --- | --- | --- |
| Live/Dead e780 |  | ThermoFisher |
| CXCR5 BV421 | J252D4 | Biolegend |
| PD-1 BV785 | EH12.2H7 | Biolegend |
| CD4 PerCP-eFluor710 | SK3 | ThermoFisher |
| CD20 B570 | 2H7 | Biolegend |
| CD25 PeCy7 | BC96 | ThermoFisher |
| OX40 PE | BER-ACT35 | Biolegend |
| Granzyme B AF647 | GB11 | Biolegend |
| Perforin FITC | B-D48 | Biolegend |

**Supplementary Table 8.**
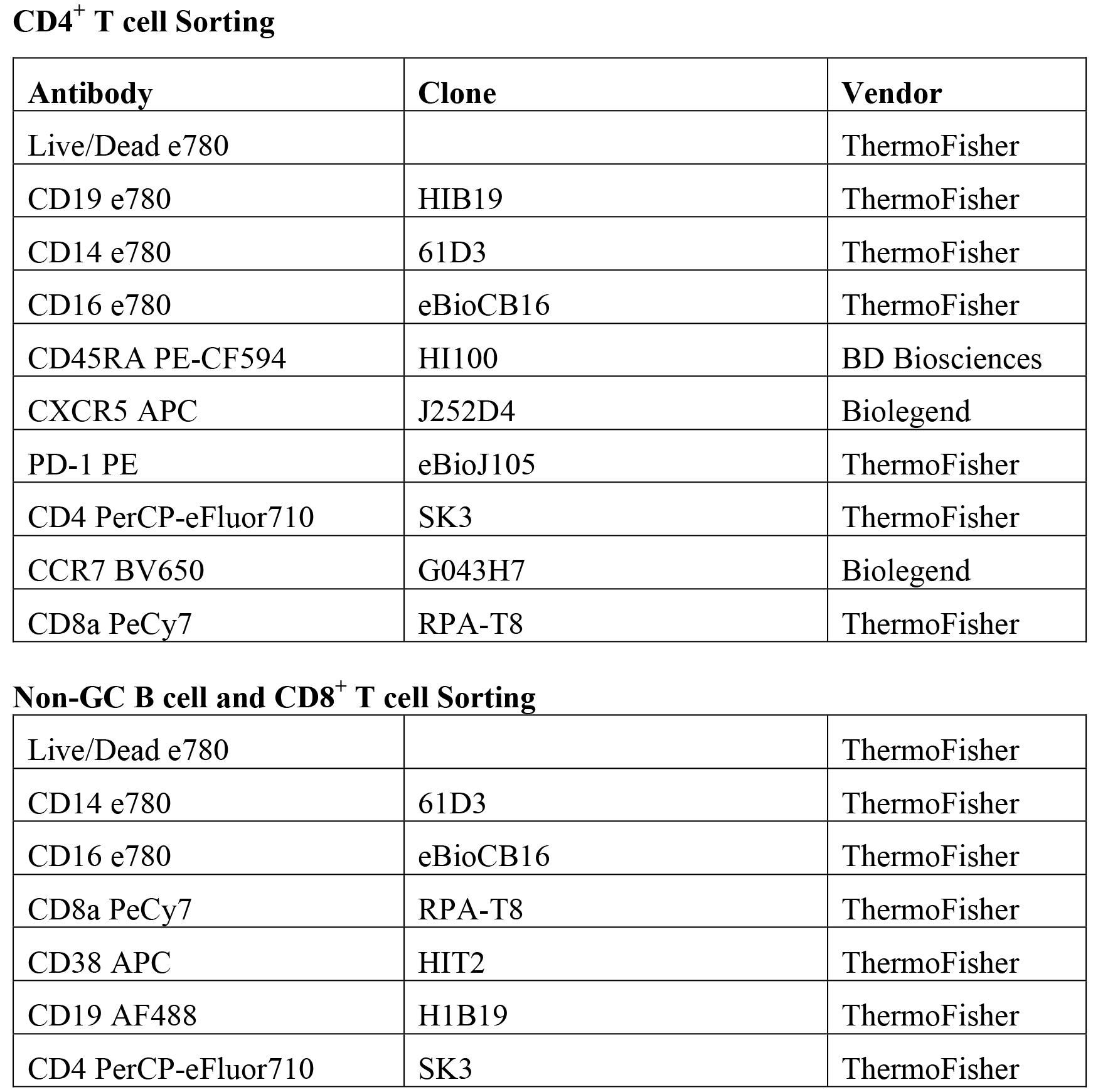
Flow cytometry antibodies for used for sorting for Cytotoxicity Assay

**CD4<sup>+</sup> T cell Sorting**
| **Antibody** | **Clone** | **Vendor** |
| --- | --- | --- |
| Live/Dead e780 |  | ThermoFisher |
| CD19 e780 | H1B19 | ThermoFisher |
| CD14 e780 | 61D3 | ThermoFisher |
| CD16 e780 | eBioCB16 | ThermoFisher |
| CD45RA PE-CF594 | HI100 | BD Biosciences |
| CXCR5 APC | J252D4 | Biolegend |
| PD-1 PE | eBioJ105 | ThermoFisher |
| CD4 PerCP-eFluor710 | SK3 | ThermoFisher |
| CCR7 BV650 | G043H7 | Biolegend |
| CD8a PeCy7 | RPA-T8 | ThermoFisher |
| **Non-GC B cell and CD8<sup>+</sup> T cell Sorting** |  |  |
| Live/Dead e780 |  | ThermoFisher |
| CD14 e780 | 61D3 | ThermoFisher |
| CD16 e780 | eBioCB16 | ThermoFisher |
| CD8a PeCy7 | RPA-T8 | ThermoFisher |
| CD38 APC | HIT2 | ThermoFisher |
| CD19 AF488 | H1B19 | ThermoFisher |
| CD4 PerCP-eFluor710 | SK3 | ThermoFisher |

